# High-throughput engineering of ligand-activated splicing ribozyme through domain insertion

**DOI:** 10.64898/2026.05.21.726912

**Authors:** August Staubus, Ella Ramamurthy, Anika Gupta, Madison Furnish, Arjun Khakhar, James Chappell

## Abstract

Domain insertion is an established method to engineer ligand-mediated control of activity in protein scaffolds. Whether this strategy can be systematically applied to large, structured RNAs remains unclear. In this study, we investigated the feasibility of engineering ligand-activated splicing ribozymes (LASRs) from group I catalytic introns. Using domain-insertion profiling coupled with high-throughput screening, we mapped the nucleotide-resolution landscape of aptamer insertion across the ribozyme and identified sites that support robust ligand-dependent control. We showed LASRs function across multiple kingdoms of life, including diverse species of bacteria and even fungi, and can be used to regulate various genetic outputs. Finally, we integrated LASRs with a genetic recorder that writes information into ribosomal RNA, enabling sequencing-based recovery of intracellular chemical signals from microbial consortia. This work establishes LASRs as an RNA-based inducible control platform for sensing diverse chemical inputs, regulating the expression of diverse genes of interest, and recording intracellular information.

## INTRODUCTION

Establishing ligand-mediated allosteric control over cellular processes is a central goal of synthetic biology^1^. For proteins, domain insertion has emerged as a powerful and generalizable strategy for converting ligand binding into regulated activity^2–7^. Inspired by how proteins naturally evolve, this approach involves embedding a sensing domain within a functional scaffold and has been used to engineer ligand sensing into diverse proteins, including GFP reporters^4^, Cas9 enzymes^5^, and metalloproteins^6^. Advances in molecular cloning and high-throughput screening have enabled systematic interrogation of allosteric potential through domain-insertion profiling across entire protein sequences, yielding generalizable design principles. Whether a similar strategy can be systematically applied to other biomolecules, such as RNA, remains unclear.

Ligand-responsive RNA switches have both been discovered in nature and engineered in laboratories and can be used to control diverse cellular phenotypes and processes, including fluorescence^8^, catalysis^9–11^, transcription and translation^12,13^, and RNA processing^14^. Synthetic RNA switches are typically constructed from relatively small scaffolds, such as translational control elements (e.g., RBSs) and self-cleaving ribozymes, to which ligand-binding aptamers are coupled, often via simple architectures such as tandem fusions^15^. Although this approach has enabled ligand-dependent control of gene expression, it has not led to a general strategy for introducing allostery into larger, highly structured RNAs. Such RNAs are involved in a wide variety of cellular processes, including splicing, translation^16^, and gene regulation^17^, but engineering such molecules present challenges due to their size, complex tertiary interactions, and conformational dynamics. As a result, RNA allostery remains poorly understood from a systematic, design-driven perspective. In this study, we ask whether high-throughput domain insertion of ligand-binding aptamers can be applied systematically to a large, structured RNA scaffold to generate robust and programmable allosteric control, focusing on the group I intron splicing ribozyme.

Group I intron splicing ribozymes are a phylogenetically widespread class of catalytic RNA that excise themselves from a transcript and ligate the flanking RNA exons^18^ (**Fig. 1AB**). While naturally occurring, these ribozymes have been extensively engineered to create genetic control elements^19^, RNA-editing technologies^20^, and mRNA biosensors^21–23^, forming the basis of biotechnologies with diverse applications that span recording gene transfer in microbial communities^24^ to the development of novel therapeutics^25,26^. A key advantage of this scaffold is its versatility. It requires only universally available cofactors (magnesium and GTP) and no protein cofactors, and thus is portable across kingdoms and can function in bacteria, plants^23^, and mammalian cells^22^. Moreover, as only a uracil is required for splicing, its exon sequences can be readily exchanged to couple splicing to diverse outputs. Despite these advantages, the potential of group I intron splicing ribozymes for programmable ligand-mediated allostery remains largely unexplored. Only a single naturally ligand-responsive group I intron has been reported^27^, and prior engineering efforts have relied on *ad hoc* aptamer insertions into a handful of sites^28^. A systematic, quantitative interrogation of allosteric potential via domain-insertion profiling has not yet been undertaken.

**Figure 1.**
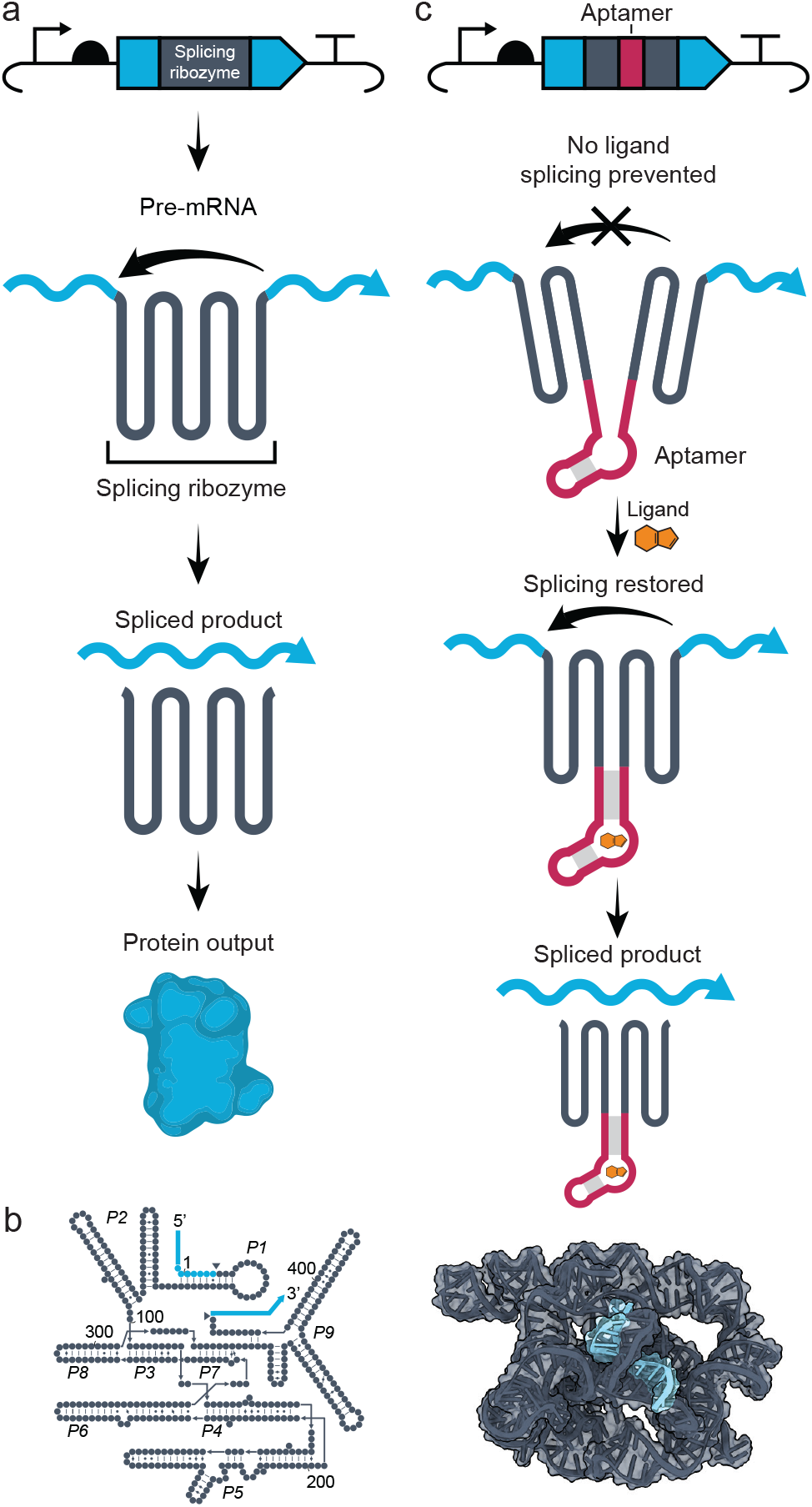
Mechanism and structure of the group I intron splicing ribozyme. **a** When transcribed from DNA, the group I intron splicing ribozyme (grey) catalyzes a splicing reaction, removing itself from the flanking exons (blue) and ligating them together. **b** The secondary (left) and tertiary (right) structure of the group I intron ribozyme, with the nucleotides of the flanking exons colored blue. **c** By inserting an aptamer into key sites of the intron, splicing becomes contingent upon the presence of a ligand.

In this study, we present the first systematic investigation of the ligand-mediated allosteric potential of a large RNA and establish splicing ribozymes as a generalizable inducible control platform, which we term ligand-activated splicing ribozymes (LASRs) (**Fig. 1C**). Using domain-insertion profiling coupled to high-throughput screening, we generated the first nucleotide-resolution maps of allosteric potential in a splicing ribozyme using multiple aptamers, revealing design principles and identifying sites that support robust ligand control. This screen resulted in a panel of LASRs with up to 20-fold activation. We show LASRs offer broad host range portability and can function across kingdoms of life (from bacteria to fungi). We then demonstrate the broad biotechnological versatility of LASRs by using them to regulate diverse genetic outputs. Finally, we integrate LASRs with a ribozyme-based genetic recorder that writes information into ribosomal RNA (rRNA), enabling autonomous recording of intracellular chemical signals in a simple synthetic microbial consortium that can be recovered by DNA sequencing.

## RESULTS

### Mapping the allosteric landscape of a splicing ribozyme to create theophylline-responsive LASRs

To map sites in a splicing ribozyme that allow for ligand-mediated allosteric control of catalytic activity when combined with a ligand-binding aptamer domain, we used saturated programmable insertion engineering (SPINE)^29^ to generate a comprehensive domain insertion library in which the theophylline-binding aptamer was inserted at all possible positions within the *Tetrahymena thermophila* group I intron (**Fig. 2A, Supplementary Fig. 1**). Randomized trinucleotide sequences were included on either side of the aptamer to serve as a communication module. These modules are often used in RNA engineering and are thought to transmit the ligand-binding state of the aptamer to the surrounding RNA scaffold. Although communication modules are widely used, their design rules are poorly understood. Consequently, identification of functional communication modules typically relies on screening of randomized pools^30,31^. The ribozyme domain insertion library was placed in codon Y66 of the sfGFP gene, a site previously identified as producing no measurable fluorescence unless splicing occurs (**Fig. 2A**)^21^. A constitutive mCherry reporter was also encoded on each plasmid, providing a normalization factor to account for sources of intrinsic noise (**Fig. 2A**)^32^. Plasmids encoding this library were then transformed into *E. coli* K-12 MG1655 and the coverage of insertion sites was determined using targeted amplicon sequencing (**Supplementary Fig. 1**).

**Figure 2.**
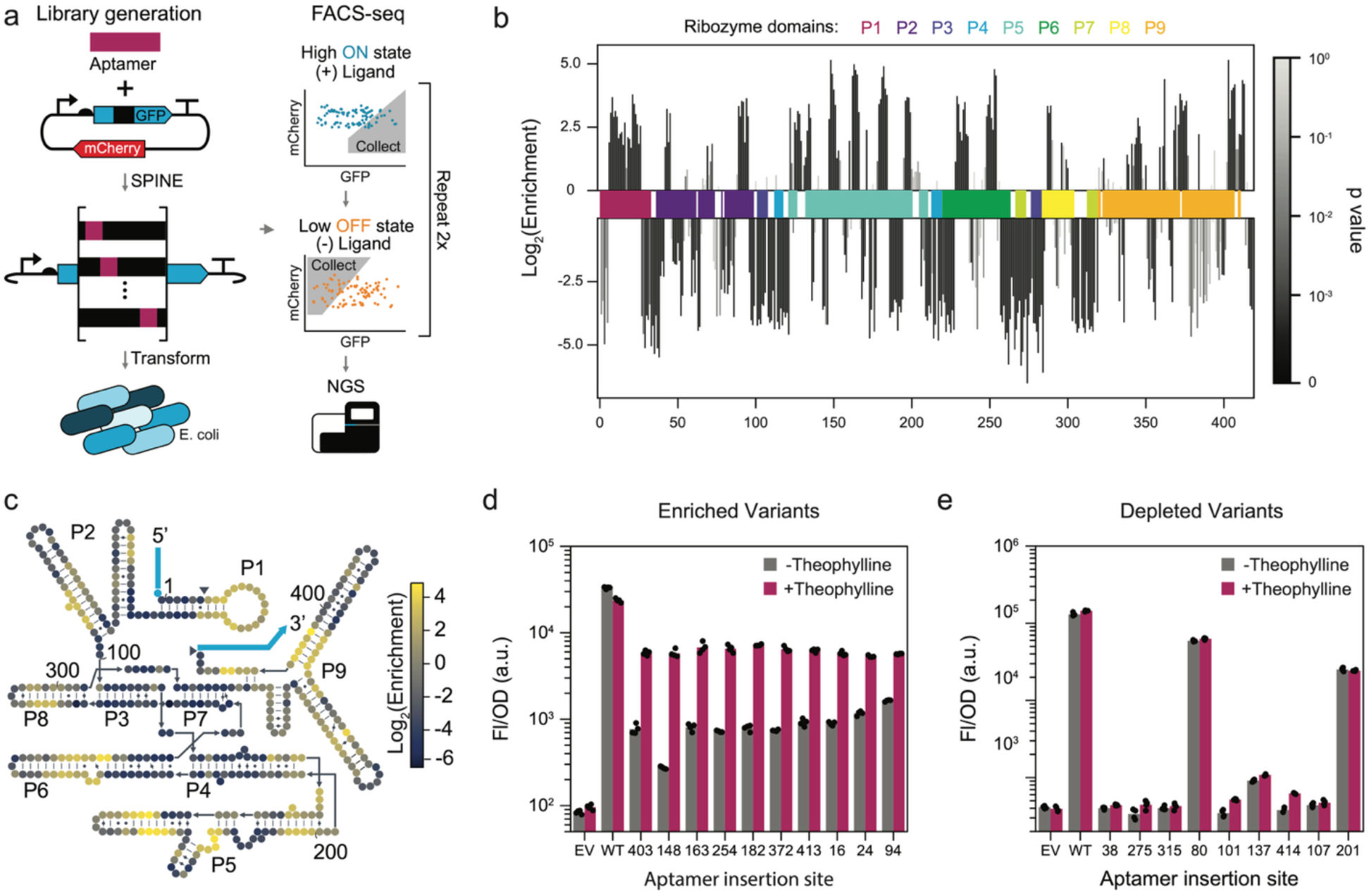
High-throughput theophylline aptamer domain insertion library screening generates ligand-activated splicing ribozymes (LASRs). **a** Schematic depicting the construction of an aptamer domain insertion library and subsequent high-throughput screening. Left, saturated programmable insertion engineering (SPINE) was used to create a comprehensive domain insertion library in which the theophylline aptamer was inserted into every possible position within the splicing ribozyme, which is embedded within a GFP reporter gene. After transforming the library into *E. coli*, cells were sorted based on the ratio of GFP:mCherry expression, with a high ratio collected after culturing in the presence of theophylline and a low ratio collected after culturing in its absence. **b** Enrichment of aptamer insertion sites after sorting. Bars represent the mean of 3 library replicates and are shaded according to p values. Ribozyme paired domains (P1-9) are indicated by colored bars. **c** Mean enrichment data overlaid onto the secondary structure of the splicing ribozyme. Fluorescence characterization (measured in units of fluorescence [FL]/optical density [OD] at 600 nm) of the splicing activity of selected (**d**) enriched and (**e**) depleted LASR designs in the presence and absence of 1mM theophylline. WT: positive control *E. coli* cells with a constitutively active wild-type ribozyme (without an aptamer). EV: negative control *E. coli* cells with an empty vector. For (**d**) and (**e**), bars indicate the mean of 4 biological replicates shown as points.

Having constructed our library, we next aimed to use fluorescent-activated cell sorting coupled with next-generation sequencing (FACS-seq) to identify library variants that were allosterically activated by theophylline (**Fig. 2A**). In brief, the transformed library was grown in the presence of theophylline and cells with a high GFP:mCherry expression ratio were collected (ON-state sort) (**Supplementary Fig. 2**). The recovered cells were then grown in the absence of theophylline, and variants with a low GFP:mCherry ratio were collected (OFF-state sort) (**Supplementary Fig. 2**). The recovered cells then underwent two additional rounds of sorting, for a total of four (ON-OFF-ON-OFF), after which the plasmids were extracted from sorted cells. Plasmids were also isolated from the naïve (unsorted) culture to quantify the initial plasmid library composition. The ribozyme region was amplified, sequenced, and analyzed to identify aptamer insertion sites and to calculate the relative enrichment of each insertion site after sorting (**Fig. 2B**).

In these data, we observed enrichment of variants with aptamer insertions located in discrete regions throughout the splicing ribozyme, suggesting that they are allosterically activated by theophylline. When overlaying the data on the ribozymes’ secondary structure (**Fig. 2C**), these regions appear to be particularly enriched within the paired (P) helical domains, particularly in domains P1, P2, P5, P6, P8, and P9. Surprisingly, a large proportion of the nucleotide insertion sites (179/418, 43%) within the ribozyme were enriched by the sorting process, suggesting that the ribozyme’s allosteric landscape is quite permissive, with multiple large and distinct regions capable of coupling ligand binding to functional activity. As expected, insertion variants within the catalytic core of the ribozyme (P3, P4, and P7) were highly depleted (mean log_2_(enrichment) −3.02) compared to the remaining domains (mean log_2_(enrichment) −0.0243), likely due to the disruption of the splicing mechanism (**Fig. 2C** and **Supplementary Fig. 3A**). Additionally, insertions into hairpin loops were depleted (mean log_2_(enrichment) −1.84) compared to insertions into paired regions (−0.529) (**Fig. 2C** and **Supplementary Fig. 3A**). This was surprising given that hairpin loops are often targets for insertions in rational RNA engineering efforts^33–41^. In contrast, some of the most enriched sites were found in the middle of helices, with paired regions on opposite sides often showing similar levels of enrichment (**Fig. 2C** and **Supplementary Fig. 3C**). Taken together, these findings unveil sequence and structural features of theophylline-response LASRs and that allosteric activation is not limited to a small set of privileged sites but instead can arise from a wide array of ribozyme regions, highlighting the unexpectedly versatile allosteric potential of this ribozyme.

We next sought to validate these findings by individually characterizing specific variants and their response to theophylline. Ten highly enriched variants were cloned, transformed into *E. coli*, and their splicing efficiency measured by GFP fluorescence in the presence and absence of 1 mM theophylline (**Fig. 2D**). Theophylline-dependent activation of splicing was observed across all enriched variants tested, ranging from 3.5 to 21-fold. Interestingly, the fluorescence level in the presence of theophylline was consistent across all measured LASR variants, whereas greater variance was seen in the absence of theophylline, with high fold-changes generally being attributable to lower off-states. As an additional confirmation of the enrichment data, we characterized a panel of highly-depleted variants and found that they had no notable response to theophylline (**Fig. 2E**). Interestingly, while most of the cloned depleted variants had little to no observable splicing activity with or without theophylline, suggesting aptamer insertion had disrupted catalytic activity entirely, two variants with aptamer insertions located within loops (site 80 and site 201) retained splicing activity, suggesting these sites tolerated aptamer insertion but were not conducive to allostery. Across all variants measured, we found that the enrichment scores correlated with the fold activation measured from the individually characterized variants (R^2^ = 0.67) (**Supplementary Fig. 4**). In contrast, enrichment correlated poorly with the ON state (+theophylline) and the OFF state (−Theophylline) fluorescence (R^2^ = 0.03 and R^2^ = 0.0 respectively), suggesting that the sorting process selected for variants with a high dynamic range, rather than a particular set level of expression. These results confirm that our domain insertion profiling is a robust approach to generating LASRs, and that enrichment via sorting is a reliable predictor of allosteric activity.

### Diversifying the inputs for LASRs

To assess the generalizability of this method for generating LASRs that respond to diverse ligands, as well as the differences in allosteric potential between different aptamers, we constructed two additional aptamer domain insertion libraries using aptamers that bind 5-aminoimidazole-4-carboxamide ribonucleotide (ZMP) and fluoride ions (**Fig. 3A**). These aptamers were selected because they are derived from the ligand-binding domains of naturally occurring riboswitches, and therefore likely exhibit substantial structural differences between their ligand-bound and unbound states^42,43^. Additionally, these ligands are chemically distinct and play different biological roles (*i*.*e*., an organic small molecule metabolite versus a negatively charged monatomic ion), allowing us to interrogate how LASRs integrate with different classes of ligands. As before, domain insertion libraries created with these two aptamers were transformed into *E. coli*, the FACS-Seq was repeated, and enrichment profiles were calculated (**Fig. 3BC**). We note one difference from the prior screen: instead of a four-sort scheme (ON-OFF-ON-OFF), a three- or two-sort scheme was used. This change was motivated by analyzing the fold enrichments from each sort of theophylline screen, which showed that additional rounds of sorting provided little benefit (**Supplementary Fig. 5**).

**Figure 3.**
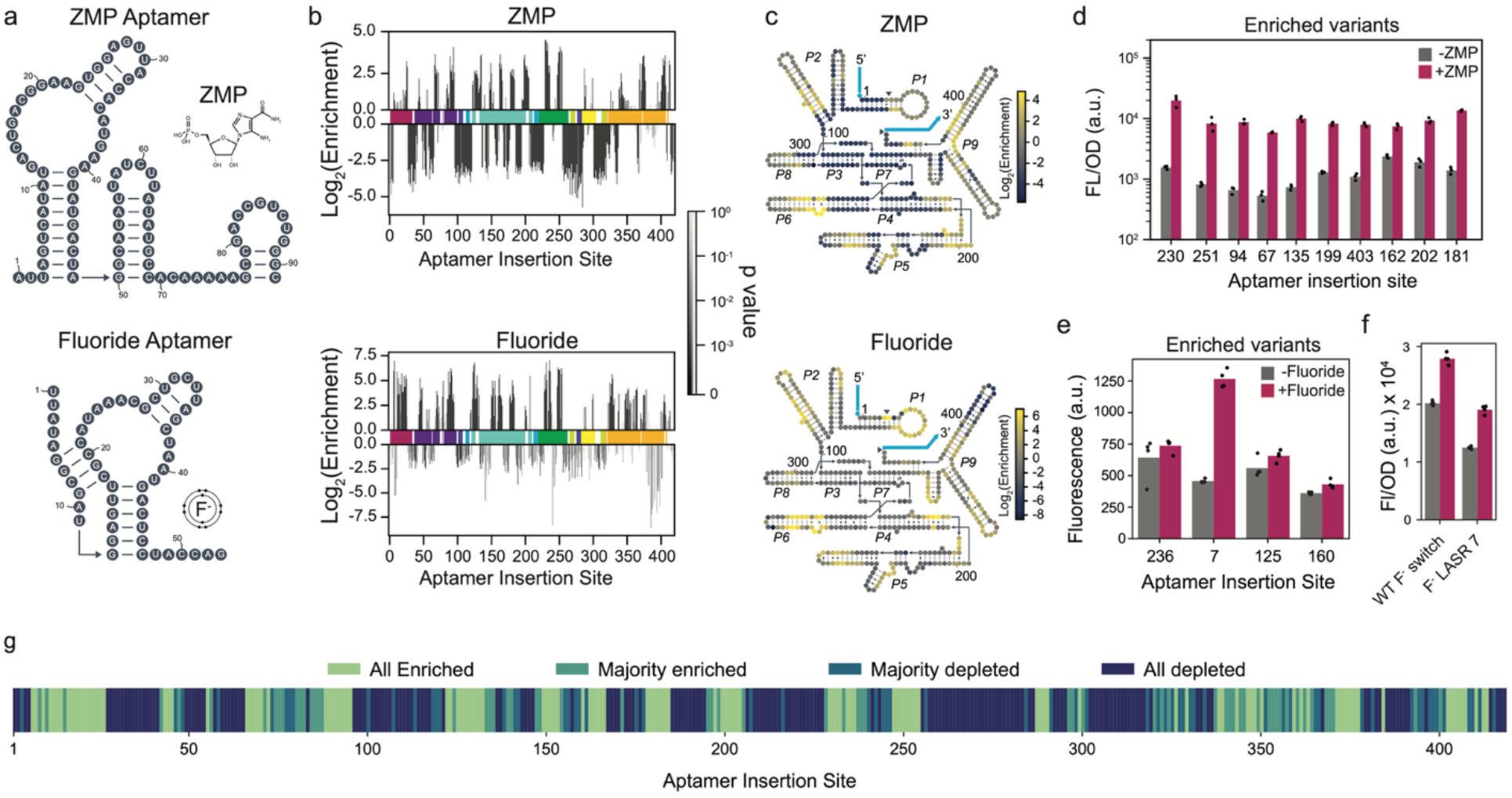
LASRs can be created using different ligand-binding aptamers. **a** Secondary structure diagrams of the ZMP (top) and fluoride (bottom) aptamers, along with their corresponding ligands. **b** Enrichment profiles for ZMP (top) and fluoride (bottom) aptamer domain insertion libraries after sorting. Bars represent the mean of 3 library replicates and are shaded according to p values. Ribozyme paired domains (P1-9) are indicated by colored bars. **c** Mean enrichment data overlaid onto the secondary structure of the splicing ribozyme. **d** Splicing activity (as measured by GFP expression) of individually cloned enriched ZMP aptamer insertion variants. Data shows bulk fluorescence characterization (measured in units of fluorescence [FL]/optical density [OD] at 600 nm) measured in the presence and absence of 1 mM ZMP. **e** Splicing activity of enriched fluoride variants and (**f**) comparison of a fluoride response LASR (at insertion site 7) to a natural fluoride riboswitch from *B. cereus*. Data shows single-cell fluorescence measured in the presence and absence of 200 µM fluoride. For (**d**) and (**e**), bars indicate the mean of 3 to 4 biological replicates shown as points. **g** Enrichment consensus at each insert site across the splicing ribozyme. Sites are color-coded according to whether they had positive enrichment values across all three aptamer domain insertion libraries (all enriched), positive enrichment in 2/3 libraries (majority enriched), negative enrichment in 2/3 libraries (majority depleted), or positive enrichment across all 3 libraries (all enriched).

The enrichment profiles for both the ZMP and fluoride aptamer libraries exhibited enrichment in a wide range of sites within the ribozyme, suggesting that the versatile allosteric potential of this ribozyme is not limited to a single aptamer-ligand pair and is instead generalizable to many other targets (**Fig. 3B**). Overlaying the enrichment data from the two new libraries onto the secondary structure of the ribozyme (**Fig. 3C**), we again observed enrichment in discrete regions across the primary sequence of the ribozyme, with insertion sites near the catalytic domains depleted and sites in structural helices enriched. To confirm that the enrichment data reflected ligand allostery, ten highly enriched and depleted variants from the ZMP and fluoride libraries were identified and individually characterized with fluorescence. For ZMP, all enriched variants showed activation in response to their respective ligands (**Fig. 3D**), while depleted variants were unresponsive (**Supplementary Fig. 6**). For fluoride, four of ten enriched variants showed a clear ligand response (**Fig. 3E**). For depleted variants, most variants showed repression in the presence of fluoride (**Supplementary Fig. 6**), suggesting that the fluoride aptamer is more disposed toward negative allostery or that fluoride exerts an indirect effect by interfering with bacterial metabolism^44^. Notably, the best-performing fluoride-responsive LASR exhibited fold activation comparable to that of the natural fluoride-responsive riboswitch (WT F^−^ switch), demonstrating that our engineered systems can match the dynamic range observed in nature (**Fig. 3F**). Taken together, these data show the generalizability of this ribozyme as a versatile allosteric scaffold across distinct aptamer inputs.

### Investigating hotspots of allostery in LASRs

We next aimed to determine whether we could uncover underlying design rules for allostery in this splicing ribozyme using our data. Allosteric hotspots – regions where ligand-binding domains are tolerated and allostery observed – have been widely observed in both natural and synthetic allosteric proteins^4,5,45,46^. We were interested in whether analogous hotspots existed in the splicing ribozyme. First, we compared the enrichment scores for each site across the three aptamer libraries (theophylline, ZMP, and fluoride). This comparison showed a strong correlation between theophylline and ZMP aptamer libraries (R^2^ = 0.67), while the theophylline-fluoride and ZMP-fluoride enrichment correlations were weaker (R^2^ = 0.29 and 0.33, respectively) (**Supplementary Fig. 7**). We then determined whether certain insertion sites were always suitable sites for creating allostery (i.e., a hotspot) and certain sites always forbidden (i.e., a coldspot) (**Fig 3G**). From this, we saw that 120 (29%) and 173 (41%) sites were enriched and depleted (respectively) across all three libraries, supporting the idea that hotspots and cold spots of allostery exist. These hotspots tended to appear in clusters within specific regions of the ribozyme, consistent with previous efforts to engineer allosteric proteins, which also found clusters of hotspots distributed throughout the primary sequence^4,5,7^. These results suggest that – like many proteins – splicing ribozymes contain clear allosteric hotspots that are compatible with a wide variety of ligand binding domains to achieve allosteric control. Moreover, they indicate that features of the scaffold may be more relevant to the allosteric function of the LASR than are features of the aptamers themselves.

To identify features of the ribozyme associated with insertion tolerance, we examined structural, sequence, and evolutionary properties of each nucleotide using a high-resolution cryo-EM structure and conservation data^47,48^. However, none of these features alone showed strong correlation with enrichment (**Supplementary Fig. 8**), and a random forest model trained on these features exhibited limited predictive power. These results suggest that insertion tolerance cannot be explained by simple local descriptors, and instead likely arises from higher-order or context-dependent interactions within the ribozyme structure.

### Interrogating the role of communication modules in LASRs

While the communication module is thought to play a key role in allosteric RNA systems, the exact function of these linking sequences remains elusive. To better understand their contribution to LASRs, we constructed libraries of randomized communication modules at two distinct aptamer insertion sites, positions 94 and 403. These sites were chosen because they showed clear switching in response to theophylline but were not responsible for the largest fold change in activation, allowing us to determine whether variance in the communication modules could improve or break function. This library was transformed into *E. coli* cells and the fluorescence distributions measured by flow cytometry (**Supplementary Fig. 9**). Compared with previously identified LASRs at these sites (termed version 1 designs), the communication module libraries exhibited a broad spectrum of activity levels, suggesting that some communication modules disrupted ligand switching while others preserved it. This finding supported a role for communication modules in governing ligand-switching behavior.

To determine which communication modules enabled ligand switching, we performed FACS-seq and targeted amplicon sequencing on the two communication module libraries. Comparing the fluorescence of the sorted population in the presence and absence of ligand revealed convergence toward a ligand-switching behavior (**Supplementary Fig. 10**) and a reduced number of variants (**Supplementary Fig. 11**), suggesting that the resulting population was enriched for functional LASRs. To identify sequence features associated with functionality, we calculated sequence conservation among the enriched variants with a log_2_(enrichment) of >2 and compared these to the naïve library (**Fig. 4B** and **Supplementary Fig. 12**). At aptamer insertion site 403, we observed a strong preference for a G at position 3, whereas at insertion site 94, there was a preference for a C at position 4. These findings indicate that functional communication modules exhibit position-specific sequence preferences that vary depending on the aptamer insertion site.

**Figure 4.**
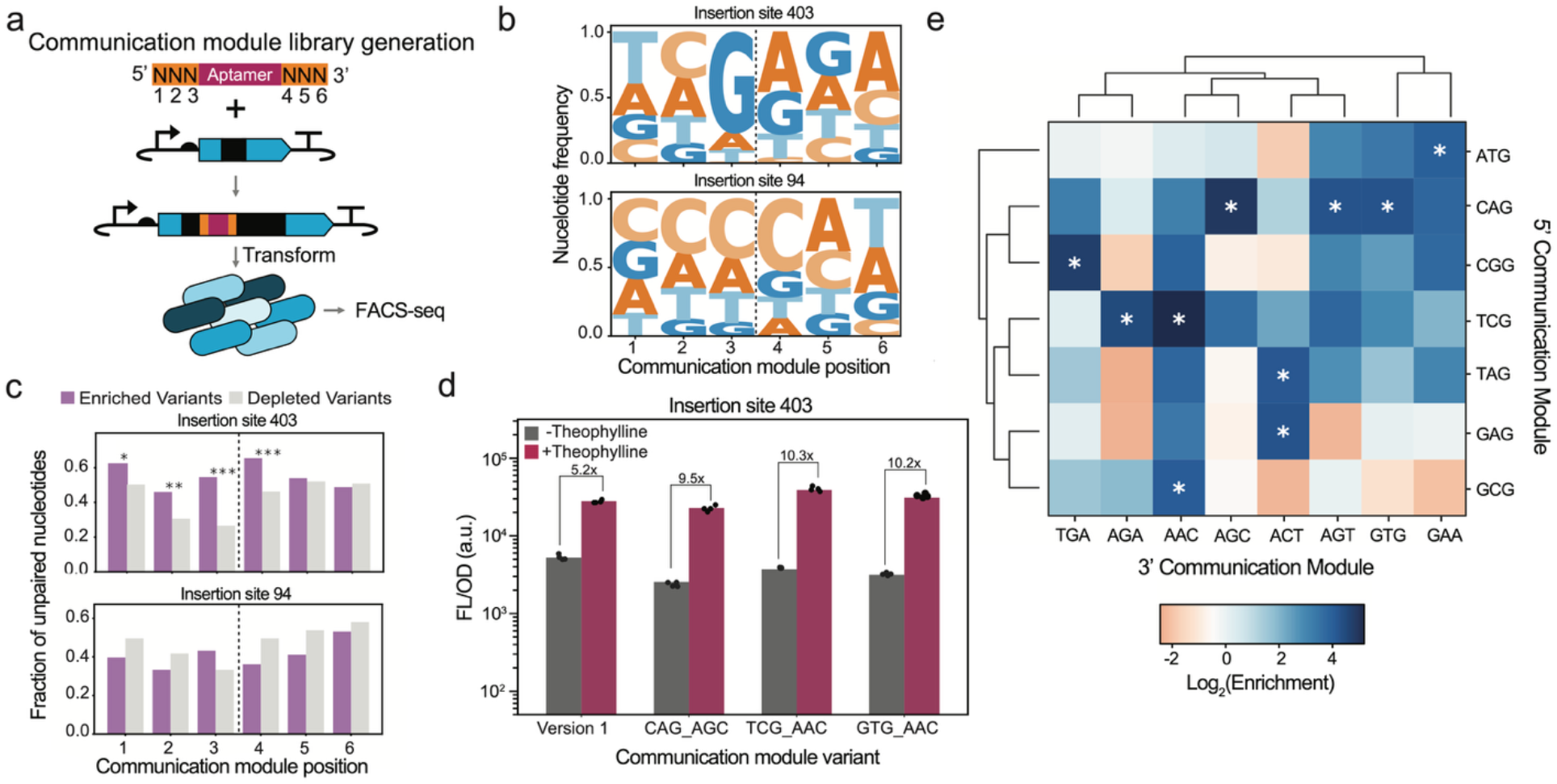
Investigating the design rules of the communication module. **a** Schematic depicting the construction and screening of a communication module library wherein an aptamer with three randomized nucleotides flanking either side (positions 1 to 6) is inserted into specific sites in the group I intron ribozyme. For each position, data were collected for 3 replicate libraries. **b** Nucleotide frequency plot of the communication modules among the enriched variants (>2 log_2_(enrichment) after FACS-seq for aptamers inserted into site 403 and 94. **c** Predicted base pairing of the communication modules for the enriched variants and depleted variants (<-6 log_2_(enrichment) after FACS-seq. Pairing data is from predicted secondary structures of the aptamer and local ribozyme sequence using NUPACK. Statistical significance was assessed independently for each position using a Fisher’s Exact Test comparing the raw counts of paired and unpaired variants within the enriched and depleted populations (*p < 0.05, **p < 0.01, ***p < 0.001). **d** Fluorescence characterization (measured in units of fluorescence [FL]/optical density [OD] at 600 nm) of the splicing activity of the three most enriched communication modules (depicted as 5’ communication module_3’ communication module) for aptamer insertion site 403 compared to the original design identified in our prior screen (version 1). Bars indicate the mean of 4 biological replicates shown as points. **e** Sequence-fitness landscapes of communication modules at site 403. The 5’ and 3’ communication modules from the top 10 most enriched variants were identified (marked by asterisks), and a combinatorial matrix was constructed displaying corresponding enrichment scores. A dendrogram showing the hierarchical clustering of the communication module sequences, grouped by their pairwise Hamming distances, is shown at the top and left of the matrix.

We next asked whether structural features were associated with functional communication modules, focusing on the base-pairing status of the communication module. Secondary structures of the aptamer and local ribozyme sequence were predicted using the NUPACK algorithm for the most enriched and depleted variants, and the pairing status of each communication module nucleotide was determined. At insertion site 403, enriched variants showed a significant preference for unpaired nucleotides within the communication module, with the strongest effect observed at positions immediately flanking the aptamer (positions 3 and 4) (**Fig. 4C**). We define unpaired nucleotides as those without a canonical or wobble base pair with the aptamer, local ribozyme sequence, or its partner communication module. In contrast, no statistically significant differences were observed at insertion site 94 (**Fig. 4C**). We further assessed whether communication modules that were unpaired with each other at specific positions were associated with higher enrichment values. Across both insertion sites, communication modules containing unpaired bases at the positions immediately flanking the aptamer (positions 3 and 4) exhibited significantly higher enrichment (**Supplementary Fig. 13**). We hypothesize that this is due to the requirement of communication modules to be flexible and respond to ligand availability; if positions closest to the aptamer are “locked” in a rigid structural interaction, they may not be able to transmit ligand binding activity to the rest of the RNA scaffold. As further validation, we constructed and individually characterized the top three communication modules from insertion sites 403 and 94, which all conformed to the sequence and structural design principles. All variants exhibited robust ligand-switching activity in the presence of 1mM theophylline, and most had greater fold change than the version 1 design (**Fig. 4D** and **Supplementary Fig. 14a**). Taken together, these results suggest that functional communication modules exhibit insertion site–dependent sequence preferences and favor an unpaired structural context at positions flanking the aptamer.

Finally, to probe the sequence–fitness landscape of communication modules, we isolated the 5′ and 3′ communication modules from the top 10 enriched variants at insertion site 403 and constructed a combinatorial matrix. The majority of pairwise combinations remained enriched, 37/56 pairs. This suggested that a functional 5’ or 3’ communication module exhibited a high degree of modularity with diverse partners. For insertion site 94, the trend was less clear, although 26/70 pairwise combinations were enriched (**Supplementary Fig. 14b**). Together, these results indicate that ligand-responsive splicing arises from the interplay between insertion site context and communication module sequence. While the insertion site largely determines whether an aptamer can be accommodated within the ribozyme, the communication module modulates the efficiency of coupling between ligand binding and catalytic activity. This division of roles motivates a two-tiered design approach, where scaffold-informed site selection is subsequently refined through sequence-level optimization of local interactions.

### LASRs are functionally portable across diverse bacterial species

After establishing and optimizing LASRs, we next wanted to understand their biotechnology potential. We first sought to understand if LASRs would function across diverse hosts. We hypothesized that LASRs would be highly portable given that group I introns are naturally found across the tree of life^49^, and that the group I intron from *T. thermophila* has been shown to function robustly across many host cell types. To test this, we evaluated the performance of LASRs in a range of species. A LASR with a theophylline aptamer at insert site 148 (LASR Theo-148) regulating sfGFP was cloned into a broad host-range plasmid (pBBR1) and transformed into a panel of gram-negative species: *Kluyvera intermedia, Hafnia alvei, Pseudomonas chlororaphis*, and *Comamonas testosteroni*. We also tested function in the gram-positive bacterium *Lactiplantibacillus plantarum* and a yeast strain *Saccharomyces cerevisiae* by using a modified version of LASR Theo-148 containing a codon-optimized sfGFP. These selected hosts span genera, families, orders, classes, and kingdoms (**Fig 5** and **Supplementary Fig. 15**). The transformed strains were then cultured in the presence and absence of theophylline, and fluorescence was measured. Across all species tested, robust activation of sfGFP expression in the presence of theophylline (ranging from 1.5 to 10-fold) was observed. These findings demonstrate that LASRs can be used reliably for ligand-inducible control of gene expression across highly diverse cell types without host-specific optimization, thereby providing a valuable new biotechnology.

**Figure 5.**
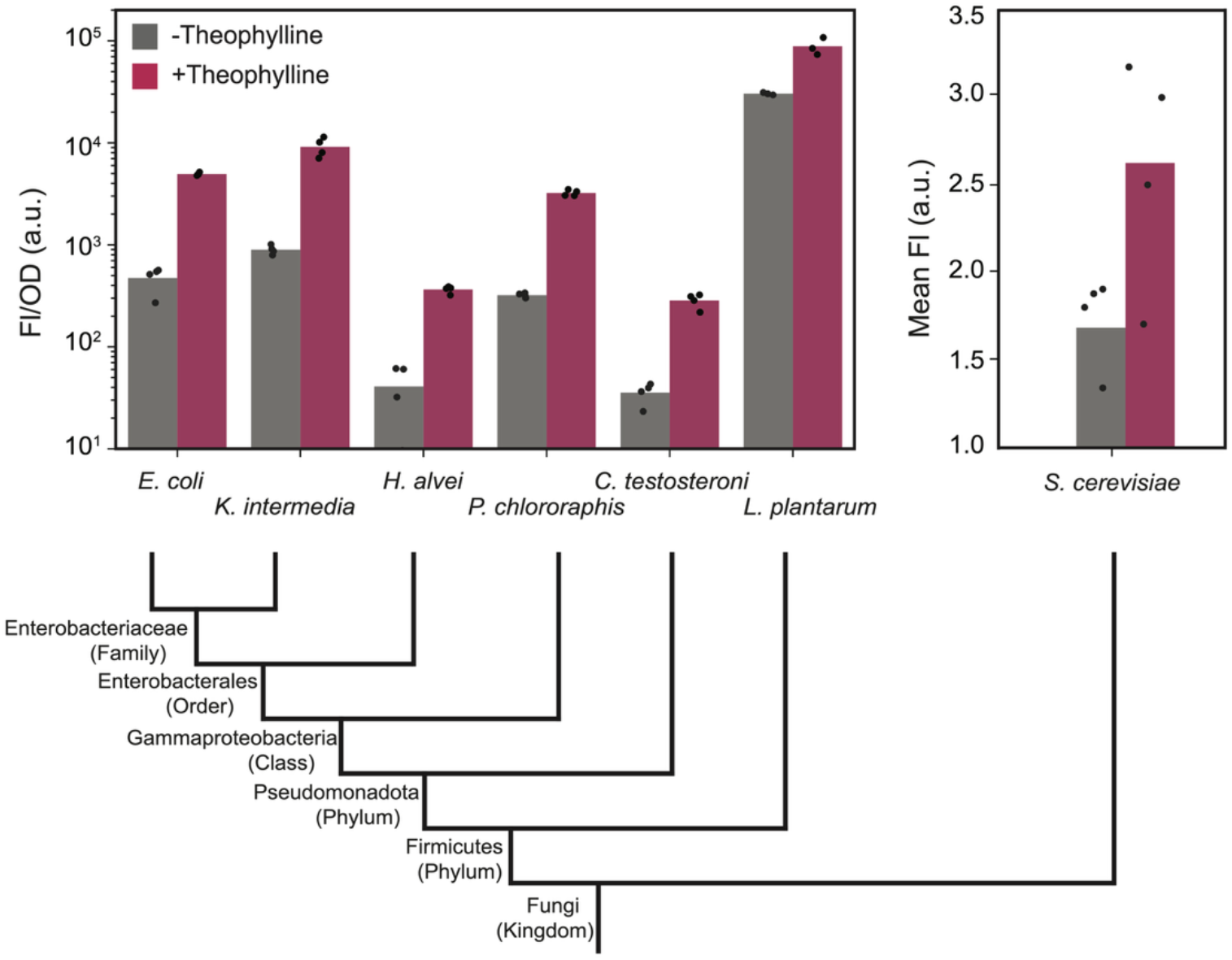
LASRs function as inducible control systems across diverse microbial species. Splicing activity (as measured by GFP expression) of LASR Theo-148 across 6 different bacterial species (*E. coli, K. intermedia, H. alvei, P. chlororaphis, C. testosteroni*, and *L. plantarum*) and a fungal species (*S. cerevisiae*). For bacterial species, data shown are bulk fluorescence characterization (measured in units of fluorescence [FL]/optical density [OD] at 600 nm), and for fungi, data shown are single-cell fluorescence by flow cytometry, both measured in the presence and absence of 1 mM theophylline. Bars indicate the mean of 4 biological replicates shown as points. The phylogenetic relationship of species is shown at the bottom of the panel.

### Creating a modular LASR genetic design for controlling different gene outputs

We next sought to create a modular design for LASRs that would allow for the facile exchange of the output gene. In our original design, the splicing ribozyme was placed within the coding sequence of the output gene (sfGFP). While this design worked well and produced minimal signal in the absence of splicing, choosing an appropriate site to insert the ribozyme inherently requires structural and functional information about the regulated gene. Additionally, as the output gene is varied, sequences within the splicing ribozyme need to be redesigned to maximize splicing activity. To create a more generalizable and modular design, we tested two new LASR architectures. In the first design, LASR Theo-148 was placed between the RBS and the start codon of the regulated coding sequence while in the second design the LASR was placed within the start codon (immediately after the ‘T’ of the ATG codon) (**Fig. 6A**). Both designs showed activation in response to theophylline, with the first design showing both a higher ON and higher OFF state than the original design and the second showing a lower ON and OFF state (**Fig. 6B**). Because the first modular design showed a larger dynamic range (comparable to that of the original design), we proceeded with this design in subsequent experiments.

**Figure 6.**
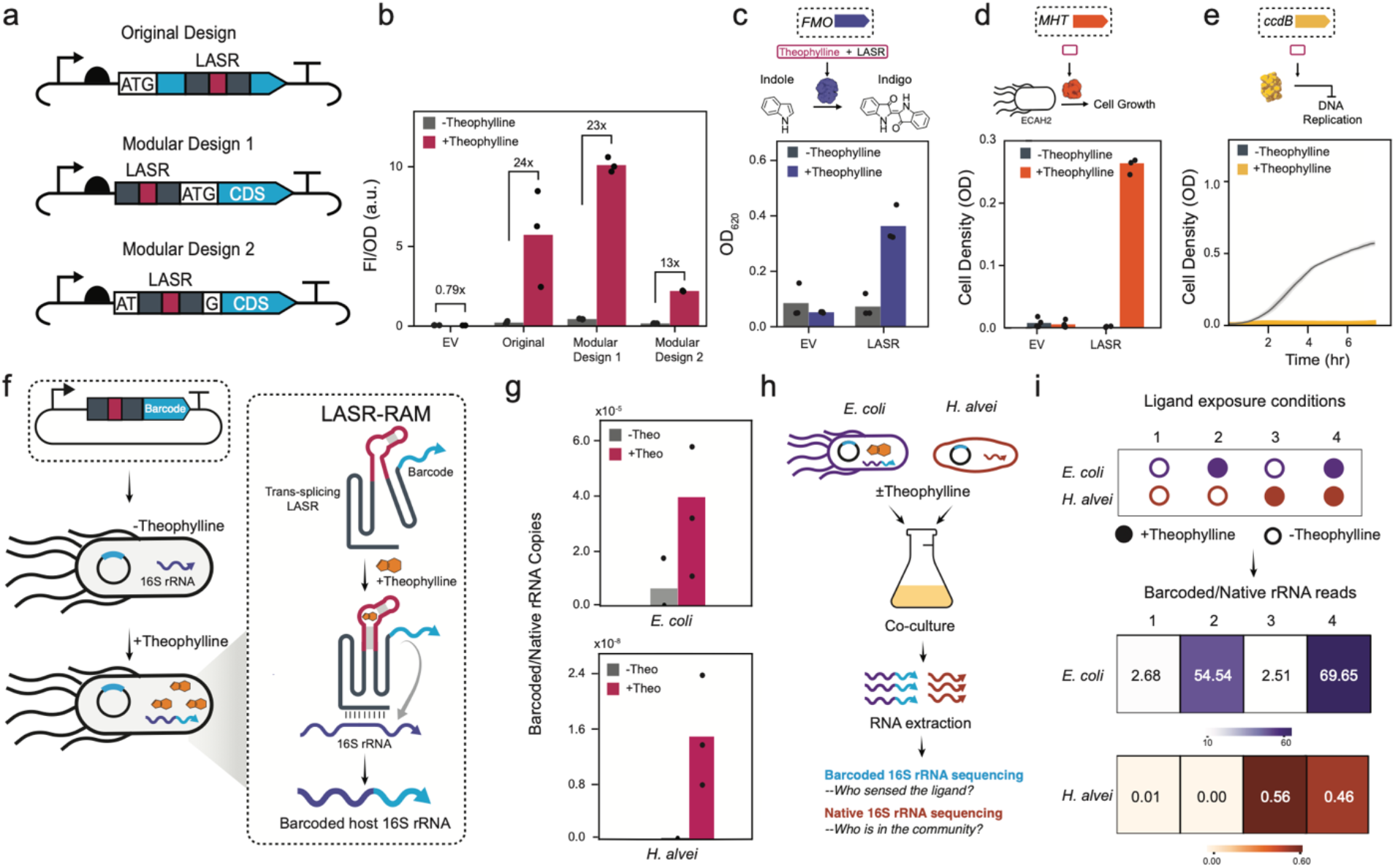
Modular LASR design enables inducible control of diverse protein outputs and the recording of intracellular chemical information. **a** Schematic comparing the original LASR design, in which the LASR was inserted in the middle of the regulated gene, and the modular designs, in which the LASR Theo-148 is either placed between the RBS and the start codon or within the start codon of the regulated gene. **b** Characterization of original and modularized LASR designs when exposed to 1mM theophylline compared to a negative control of *E. coli* cells with an empty vector. **c** Characterization of the modular LASR design using the expression of flavin monooxygenase (FMO) measured by optical density at 620 nm (OD620). **d** Characterization of the modular LASR design using the expression of methyl halide transferase (MHT) that complements auxotrophy in *E. coli* strain ECAH2 measured by optical density at 600 nm (OD600). **e** Characterization of the expression of ccdB, a toxin gene that inhibits DNA replication, measured by cell growth (OD600). **f** A schematic of the LASR-RAM system designed to splice synthetic RNA barcodes onto host 16S rRNA only when theophylline is present. **g** Quantification of native and barcoded 16S rRNA in the LASR-RAM system in *E. coli* and *H. alvei* in the presence and absence of theophylline using RT–qPCR. **h** A schematic of a consortium experiment in which *E. coli* and *H. alvei* containing LASR-RAM are cultured independently in the presence or absence of theophylline and then combined at an equal ratio. Barcoded and native 16S rRNA is then extracted and sequenced to determine who was exposed to theophylline (barcoded 16S rRNA) and the community composition (native 16S rRNA). The top of the panel shows ligand exposure conditions, and the bottom of the panel shows the ratio of barcoded and native 16S rRNA reads for *E. coli* and *H. alvei*. For (**b**), (**c**), (**d**), (**e**), (**g**) and (**i**) bars indicate the mean of 3 or 4 biological replicates shown as points.

To evaluate whether our modular design could be used to regulate the expression of different output genes, we combined LASR Theo-148 with output genes of diverse functions. First, we evaluated the regulation of the flavin-containing monooxygenase (FMO) gene from *Methylophaga aminisulfidivorans st. MP*^*33*^ that catalyzes the conversion of tryptophan-derived indole into indoxyl, which in turn forms indigo, a bright blue pigment. Characterization of this design in *E. coli* revealed tight control of indigo production in the absence of ligand, and strong production in its presence (**Figure 5C**). Next, we measured the ability of the modular LASR to regulate the expression of methyl-halide transferase (MHT), a gas-producing reporter used for measuring gene expression in environments like soil^50,51^. To do this, we used a growth-complementation assay in the *E. coli* strain ECAH2. When grown under cysteine-limiting conditions, this strain can only use S-adenosylmethionine (SAM) as a sulfur source if it expresses an active MHT and NaBr is present^52^. We observed that ECAH2 cells transformed with the MHT-LASR construct grew robustly only in the presence of theophylline (**Fig. 5D**). Finally, we evaluated the regulation of the toxin gene *ccdB*, which inhibits DNA gyrase and induces cell death. Measuring the growth rate of *E. coli* transformed with ccdB-LASR plasmid, we saw a significant decrease in growth rate in the presence of theophylline (**Fig. 5E** and **Supplementary Fig. 16**). Taken together, these results show that we can establish a modular LASR design and are able to achieve robust control of diverse gene outputs, including metabolic enzymes, toxins, and reporters.

### Coupling LASR to a genetic recorder enables the recording of intracellular chemical information in simple microbial consortia

We next sought to determine whether LASRs could be coupled to a genetic recorder to record the presence of a ligand in the intracellular environment. Recently, we showed that a *trans*-splicing ribozyme could be engineered to barcode 16S ribosomal RNA (rRNA). We showed that when this genetic reporter, called RNA-addressable modification (RAM), is embedded into mobile DNA, it can be used to barcode host cell rRNA to record gene transfer^24^. Barcoded rRNA can then be isolated, amplified, sequenced, and analyzed to identify which members of a microbial community participated in gene transfer. Given that RAM and LASRs are derived from the same ribozyme, we hypothesized that it would be possible to create a ligand-responsive RAM (LASR-RAM) that barcodes host-cell 16S rRNA in the presence of a ligand.

To test this hypothesis, we cloned the aptamer and communication module from a theophylline-responsive LASR (Theo-148) into the equivalent site in RAM. In the presence of theophylline, this LASR-RAM system is designed to amend an RNA barcode onto the host 16S rRNA, forming a transient record of the ligand’s presence (**Figure 6F**). To evaluate this function, we transformed a plasmid encoding this design into *E. coli* cells and grew them in the presence or absence of theophylline. We then extracted the RNA from these samples and quantified the barcoded and unbarcoded (native) 16S rRNA by RT-qPCR. Comparing the barcoded 16S rRNA to native 16S rRNA abundance revealed a strong increase in barcoded 16S rRNA, confirming ligand-dependent activation of RAM (**Figure 6G**). To determine whether LASR-RAM can efficiently record theophylline presence in different microorganisms, we cloned it into a broad-host-range plasmid (pBBR1) and transformed this plasmid into *H. alvei*. In the presence of theophylline, *H. alvei* showed barcoding efficiencies comparable to those of *E. coli* (**Figure 6G**). These findings showed that LASRs could be effectively coupled with RAM to establish ligand-responsive rRNA barcoding.

We next investigated whether we could use LASR-RAM to demultiplex chemical information in a simple microbial community to ask which strains contained an intracellular chemical. To do so, cultures of *E. coli* and *H. alvei* transformed with LASR-RAM were grown either in the presence or absence of theophylline, washed, and then combined into a synthetic community. We also spiked in a fixed number of *P. chlorophasis* cells expressing a constitutively active (*i*.*e*., not ligand-responsive) version of RAM. Total RNA was then extracted, barcoded and native 16S rRNA sequenced, and the read abundances of *E. coli* and *H. alvei* were normalized to the *P. chlorophasis* reference (**Figure 6H**). Analysis of native rRNAs revealed that in the absence of theophylline, *E. coli* and *H. alvei* represent equal portions of the total amplicon sequence variants (ASVs) as expected (**Supplementary Fig. 17)**. Analysis of the barcoded 16S rRNA showed relative increases in RAM signal for *E. coli* and *H. alvei* corresponding strongly with their ligand exposure condition. Taken together, this shows that LASR-RAM can be used to effectively record intracellular chemical information in a synthetic microbial consortium that can be easily read out using amplicon sequencing.

## DISCUSSION

This work represents the first systematic domain insertion profile study of ligand-responsive allostery in a large RNA. By performing this research, we discovered that the splicing ribozyme is highly permissive for allostery, with many insertion sites capable of yielding functional LASRs. We propose that this scaffold offers a substantial advantage over traditional small scaffolds (e.g., RBSs and terminators), and that its large sequence space and complex secondary and tertiary structure provide a broad design space for engineering allostery. Although we found it difficult to identify nucleotide features that correlated with allostery, we posit that, with further studies across different scaffolds, it should be possible to uncover the design principles that enable more predictable design. Additionally, the molecular mechanism of signal transduction – how ligand-induced structural changes in the aptamer propagate through the ribozyme scaffold – remains unclear. In the future, leveraging RNA chemical structural probing technologies could provide a tractable method to begin to answer this question^53^.

Despite demonstrating that this form of allostery can be readily engineered, such ligand-controlled ribozymes are surprisingly rare in nature, with only a single natural ligand-responsive splicing ribozyme identified to date^27^. The RNA world hypothesis posits that early life relied on RNA for both catalysis and regulation, which would seem to favor the evolution of such systems. Yet ligand-mediated allostery in ribozymes is largely absent from modern biology^54^. Here, we show for the first time that establishing allostery through domain insertion is surprisingly straightforward in a large RNA such as a splicing ribozyme. Domain insertion is a dominant form of allostery evolution in proteins, so its scarcity in modern allosteric RNAs might reflect other factors, such as reduced genetic stability or competition with protein-based regulatory systems^54^.

Ligand-responsive RNA switches have long been of interest for biotechnology^55^. In large part, this is because new RNA aptamers can be easily generated through *in vitro* selection (i.e., SELEX^56,57^) or, more recently, de novo using AI^58,59^, opening the possibility for custom switch design against user-defined inputs. Compared to prior scaffolds, we believe splicing ribozymes offer several advantages for the generation of new sensors. First, our study clearly shows a surprisingly large design space for creating allostery in splicing ribozymes, indicating they may be a particularly versatile platform for finding allosteric solutions with novel aptamers. Second, unlike gene regulatory elements like RBS or terminators, which have host-specific design rules and do not readily translate across different hosts, LASRs can function across diverse bacteria and fungi using a single design. While the dynamic range of these systems could likely be further enhanced through host-specific optimization, this result nonetheless highlights the broad portability and utility of LASRs. Thus, LASRs add to a growing toolbox of broad host range genetic parts^60–62^, which is of increasing interest as non-model and environmental microbes are being explored for industrial and environmental applications.

In addition to complementing existing ligand-sensing switches, LASRs also enable new applications. To date, most efforts have been placed upon engineering of ligand-response proteins such as transcription factors and two-component sensors^63,64^. Like protein-based switches, LASRs offer output modularity and can be diversified by altering the sensing component (e.g., exchanging aptamers). Additionally, because LASRs are a single-RNA system, they offer a broad host range and a minimal genetic footprint. For example, establishing ligand control over a native gene would only require insertion of the LASR sequence into the native coding sequence, without the need for additional regulatory elements (e.g., promoters, terminators, or RBS). We anticipate that these features will be particularly useful for the engineering of non-model species and microbial communities. Leveraging these benefits, we also demonstrate novel applications, including coupling LASR to RNA-Addressable Modification (RAM) to enable the recording of intracellular chemical events within ubiquitous 16S rRNA sequences across a simple microbial consortium. In the future, we anticipate this could be useful for a variety of applications, including mining membrane transporters, identifying potential host cells suitable for bioremediation of human-made pollutants, and biosensing. We also anticipate that LASRs could be coupled with DNA genetic recorders for permanent genetic recordings^65^.

All together, these findings provide new fundamental insights into the allosteric capacity of large, enzymatic RNAs while simultaneously establishing their utility as a modular and broad host range biotechnology, with applications ranging from inducible control to genetic recordings.

## ACKNOWLEDGEMENTS

This work was supported by The Welch Foundation grant C-2183-20240404 J.C. and the National Science Foundation CAREER grant no. 2237512, to J.C. M.F. and A.K. were supported by a grant from NIGMS (5R35GM155313).

## CONTRIBUTIONS

J.C. and A.S. conceived the study. Experiments were performed by A.S, E.R, A.G, and M.F. A.S., E.R, and J.C. wrote the paper, and A.G, M.F, and A.K provided feedback.

## DATA AVAILABILITY

Source data is provided with this paper.

## METHODS

### Plasmid assembly, strains, and antibiotic selection

All plasmids used in this study are listed in **Supplementary table 1**. The key plasmids are visualized in **Supplementary table 2**. All assembled plasmids were verified using Sanger DNA sequencing or Nanopore sequencing. Aptamer sequences used are shown in **Supplementary Table 3**. Strains used in this study are shown in **Supplementary table 4**. For antibiotic selection, kanamycin (100 µg/mL for *E. coli* and 200 µg/mL for all other gram-negative strains), erythromycin (10 µg/mL), and spectinomycin (100 µg/mL) were used.

### Insertion library cloning

The aptamer insertion libraries were cloned using the SPINE method, described previously^29^. First, a staging library was cloned in which Golden Gate restriction enzyme sites were introduced into every position. To do this, the sequence of the *Tetrahymena thermophila* group I intron was computationally divided into three ‘tiles’ each comprising one-third of the intron. Oligos were designed that were identical to the first tile, with the exception that a genetic handle (two outward-facing BsaI restriction sites) was placed between the first and second nucleotides in the first oligo, between the second and third nucleotides in the second oligo, and so on. Each oligo was flanked by two BsmBI restriction sites, which in turn were flanked by two primer binding sites specific to each tile. This process was repeated for the remaining two-tiles of the intron sequence, resulting in three total oligo pools.

Pooled oligos were synthesized and amplified by PCR using primers that bound to the tile-specific primer binding sites and gel extracted. Three plasmids encoding a ribozyme inserted into sfGFP were cloned, in which the wild-type sequence corresponding to each tile was replaced with BsmBI restriction sites. Amplified tiles were then cloned into each corresponding plasmid using Golden Gate cloning and transformed by heat shock into *E. coli* MG1655 cells. Serial dilutions of the transformation were performed to assess transformation efficiency and ensure adequate library coverage. Recovered cells were grown overnight in 10 mL LB and antibiotic (LB-Ab) at 30°C with shaking at 225 rpm. Glycerol stocks were made by combining overnight cultures with an equal volume of sterile 50% glycerol and stored at −80°C. Diversity of the three staging plasmids was confirmed by amplifying the ribozyme region and performing amplicon sequencing using paired-end short-read sequencing (Illumina).

Following the staging library, aptamer insertion libraries were then generated by cloning aptamers into the BsaI sites. Aptamers, randomized communication modules, and flanking BasI sites were cloned into a plasmid. To assemble each aptamer insertion library, 200 fmol of the staging library and 1 pmol of aptamer part plasmid were combined in 100 µL Golden Gate assembly reaction. Golden Gates were incubated overnight at 37 °C, then purified using a DNA clean and concentrate kit (Zymo), eluting in 10 µL of H_2_O. Eluted assemblies were divided into thirds, with each third transformed into 50 µL of MG1655 competent cells. After recovery, samples of each transformation were serially diluted and plated as described above. Three transformations were combined to 50 mL LB-Ab, grown overnight at 37°C, shaking at 225 rpm.

### Communication module library cloning

Using second-strand synthesis, double-stranded aptamer sequences were generated with degenerate nucleotides in the 6 communication module positions. These inserts were assembled into the corresponding LASR designs using Golden Gate cloning. Assemblies were performed as triplicates and transformed as described above.

### Fluorescence-activated cell sorting of libraries

100 µL of the glycerol stock of each aptamer insertion library was used to inoculate 5 mL of LB-Ab and incubated overnight at 37°C, shaking at 225 rpm. In parallel, cells containing control plasmids were also grown. After growth, cells were diluted 1:50 in fresh LB-Ab in a 24-well block. One replicate of each tile was combined at this step (3 total) proportional to their OD600, so that a single culture contained all aptamer insertions. Cultures were then incubated at 37°C, shaking at 800 rpm, until they reached mid-exponential phase (3-4 hours). They were then diluted 1:50 in fresh LB-Ab with the corresponding ligand and grown until mid-log phase. Cells were then diluted 1:200 in sterile PBS and analyzed on a Sony SH800 fluorescence-activated cell-sorter (FACS). For control samples, 100,000 events were measured. For libraries, a gate was set to select for variants with a high ratio of GFP:mCherry and 5 x 10^6^ cells were collected. After sorting, cells in the sheath fluid were transferred to an equal volume of double-concentrated LB without antibiotics and grown for 1 hour at 37 °C, shaking at 250 rpm. After recovery, antibiotics were added to the cultures, which were then returned to the incubator overnight. The following day, the same process was repeated, omitting the ligand from the second dilution step. When sorting, cells with a low ratio of GFP:mCherry were collected. The following day(s), additional on and off sorts were performed, depending on the specific library. For sorting the communication module library, cells were prepared as described above and 40,000 cells were collected each day.

### Amplicon sequencing and analysis of the insertion library

As the length of the DNA encoding the ribozyme with an aptamer inserted is >500 bp, the aptamer insertion libraries were amplified as two separate amplicons, each 250 bp. PCR primers (**Supplementary Table 5**) were appended with partial Illumina adapters using a 15-cycle PCR reaction, and the correct product was gel extracted. Sequencing was performed through Genewiz’s Amplicon-EZ service, which is performed on an Illumina MiSeq. Sequencing data was analyzed using a custom pipeline. Briefly, cutadapt was used as a quality control step to remove any extraneous sequences that lacked the primer-binding sites. Next, forward and reverse reads were merged using cutadapt. Reads were then imported into Python and dereplicated. Aptamer position was determined by aligning the aptamer sequence to each unique read. Reads with a low-quality alignment to the aptamer or that were not wild-type ribozyme were discarded. Communication modules were identified as the three nucleotides up and downstream of the aptamer. For some analyses, the data was aggregated by insert site, ignoring the communication module.

### RNA structure predictions

Nucleic Acids Package (NUPACK)^66^ was used to predict RNA secondary structures and custom python scripts used to analyze base pairing status.

### Bacterial transformations

Bacterial species used to test the portability of LASRs were transformed by electroporation. To prepare electrocompetent samples of the gram-negative strains, 500 µL LB-Ab was inoculated from a glycerol stock of each strain. Inoculated cultures were grown overnight at 30°C, shaking at 800 rpm. After overnight growth, this was used to inoculate 50 mL of the appropriate media, then cultured to an OD600 of 0.4-0.6. Cultures were washed four times with cold 10% glycerol and concentrated to a final volume of 200 µL. For electroporation, 1.5 µL of DNA (~50 ng) was added to 40 µL of electrocompetent cells. Cells were electroporated using the appropriate protocol (**Supplementary Table 6**) and then recovered in 1 mL of SOC media for 1 hour, after which they were plated on LB-AB. To prepare electrocompetent samples of *L. plantarum*, 3 mL of MRS media was inoculated from a glycerol stock of the strain. Inoculated cultures were grown overnight at 37°C, shaking at 800 rpm. After overnight growth, 300 µL of the cultures were used to inoculate 50 mL of SGMRS media, then cultured to an OD600 of 0.4-0.6. Cultures were washed two times with cold SM buffer and concentrated to a final volume of 160 µL. For electroporation, 500 ng of DNA was added to 80 µL of electrocompetent cells. Cells were electroporated using the appropriate protocol (**Supplementary Table 6**), then recovered on ice for 1 hour. Then, cells are resuspended in 1 mL of SGMRS media and recovered for 3-4 hours at 30°C, without shaking. After recovery, cells are plated on MRS media. Bulk fluorescence measurements were performed.

### Bulk cell fluorescence measurements

Individual colonies transformed with a plasmid were picked from an agar plate and used to inoculate 300 µL of LB-Ab. Cultures were incubated overnight at 37°C, shaking at 1000 rpm. The following morning, cultures were diluted 1:50 in 200 µL of fresh LB-Ab into a sterile 96-well block and incubated at 37°C, shaking at 1000 rpm until mid-log phase (3-4 hours). Cultures were then diluted 1:50 (200 µl final volume) in fresh LB-Ab media with ligand or no ligand control (e.g., DMSO) and grown until mid-log phase. After growth 50 µL was transferred to a 96 well plate and diluted 1:1 with PBS. OD600 and GFP fluorescence were measured (excitation: 485 nm, emission 510 nm) using a Tecan Spark plate reader. OD600 and GFP measurements had the values of the media subtracted.

### Yeast transformations

To prepare chemically competent samples of *S. cerevisiae*, 5 mL of 2x YPAD media was inoculated from a single colony. Inoculated cultures were grown overnight at 30°C, shaking at 225 rpm. After overnight growth, this was used to inoculate 50 mL of 2x YPAD media, then cultured to an OD600 of 0.3-0.6. After this, cells were pelleted, washed with sterile water, and concentrated to a final volume of 100 µL. Cells were transformed using a Lithium Acetate transformation method^63^, plated on uracil dropout media, and grown at 30°C for 48-72 hours. Colonies were validated using colony PCR.

### Yeast cell fluorescence measurements

Individual colonies verified to contain the plasmid were used to inoculate 4 mL of synthetic complete media (SCM). Cultures were grown overnight at 30°C, shaking at 225 rpm. The following morning, cultures were diluted to 100 events/µL in 4 mL of SCM. 1 mM theophylline (induced) or sterile water (uninduced) was added to the cultures. GFP fluorescence was measured using a BD Accuri6 Plus flow cytometer with attached CSampler. 10,000 events were collected for analysis.

### Modular LASR output assays

#### LASR-FMO

Individual colonies transformed with a plasmid were picked from an agar plate and used to inoculate 300 µL of LB-Ab. Cultures were incubated overnight at 37°C, shaking at 1000 rpm. The following morning, cultures were diluted 1:50 in 5 mL of fresh LB-Ab supplemented with 5 mM tryptophan and 1 mM theophylline (induced) or 0.5% DMSO (uninduced) and incubated at 37°C, shaking at 225 rpm for 24 hours. Indigo pigment was extracted by pelleting cells, removing supernatant, and vortexing cells in 1 mL of DMSO. Cells were centrifuged at 13,000xg for 20 minutes, and 100 µL of supernatant was transferred to a 96-well plate, and the relative indigo concentration was evaluated by measuring optical density at 620 nm using a Tecan Spark plate reader.

#### LASR-MHT

Individual colonies transformed with a plasmid were picked from an agar plate and used to inoculate 6 mL of M63 media supplemented with antibiotics and 1 mM theophylline (induced) or 0.5% DMSO (uninduced). Cultures were incubated at 37°C, shaking at 225 rpm, with OD600 measurements taken every 24 hours over 3 days to track cell growth. For each OD600 measurement, 100 µL of cells were transferred to a 96-well plate, and the optical density at 600 nm was measured using a Tecan Spark plate reader.

#### LASR-ccdB

Individual colonies transformed with a plasmid were picked from an agar plate and used to inoculate 300 µL of LB-ab. Cultures were incubated for 4 hours at 37°C, shaking at 1000 rpm. After 4 hours, cultures were diluted 1 µL of cells in 99 µL of LB supplemented with antibiotic and 1 mM theophylline (induced) or 0.5% DMSO (uninduced) in a 96-well plate. Cell growth was recorded by measuring optical density at 600 nm every 10 minutes over 8 hours in a Tecan Spark plate reader.

### RT-qPCR

Cultures were prepared for *H. alvei* and *E. coli* MG1655 as described for bulk cell fluorescence measurements. Following incubation, RNA was extracted using bead beating (with 3 x 2-minute pulses with 1-minute ice rests) and RNA was extracted using the Zymo Quick-RNA Extraction Kit. RT-qPCR was performed as described in Kalvapalle et al. *Nature Biotechnology* (2025)^24^.

### LASR-RAM microbial consortia measurements

Bacterial strains *H. alvei* and *E. coli* MG1655 were transformed with the LASR-RAM plasmid and *P. chlororaphis* with the constitutive RAM plasmid via electroporation, as described above. *P. chlororaphis* was included as a reference strain to normalize *H. alvei* and *E. coli* relative read frequency. All strains were grown as described for bulk cell fluorescence measurements, and the *H. alvei* and *E. coli* cells were induced with 1mM theophylline. Cultures of *E. coli* and *H. alvei* were combined in proportion to their OD600 and *P. chlorophasis* was added at 10% of the total number of cells, as a reference strain, for a total volume of 1 mL LB-Ab. After combining, cultures were pelleted, lysed via bead beating, and RNA extracted using the Zymo Quick-RNA Extraction Kit. To convert extracted RNA to DNA for amplicon sequencing, the RNA was reverse-transcribed into cDNA using SuperScript IV and a blocking oligo to prevent off-target amplification. Two rounds of PCR amplification were then performed to amplify the barcoded product and add Illumina adaptor sequences. PCR products were sequenced by Genewiz using their Amplicon-EZ protocol. The LASR-RAM NGS data was analyzed using a previously described protocol in Kalvapalle et al. *Nature Biotechnology* (2025)^24^.

