## Supplementary Figures and Tables for "High-throughput engineering of ligand-activated splicing ribozyme through domain insertion"

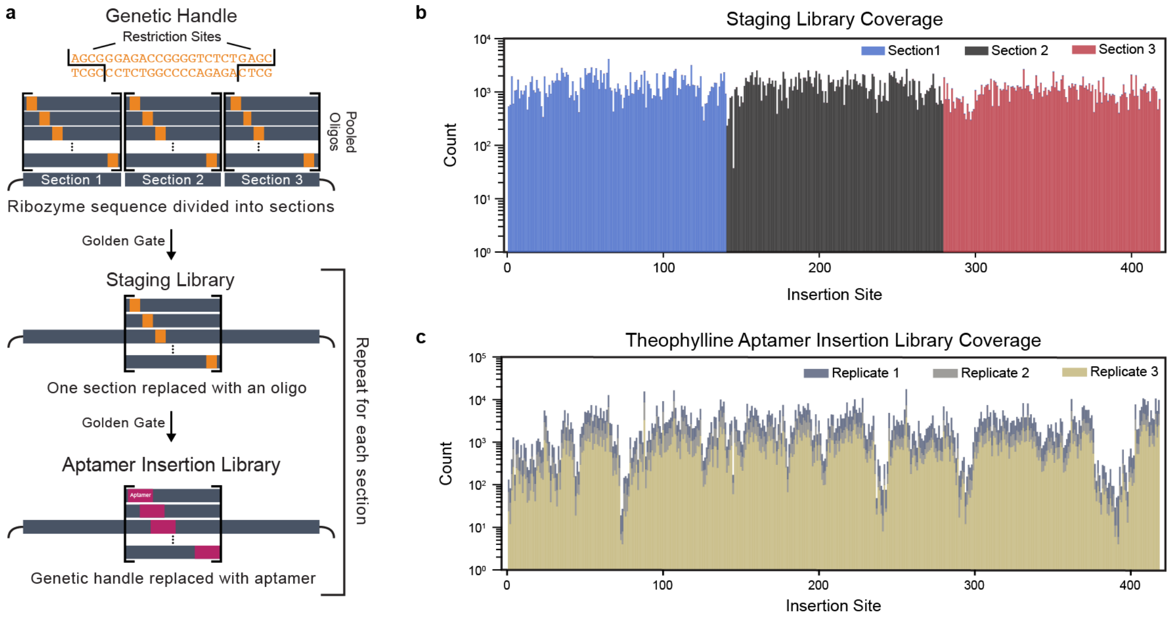

**Supplementary Fig. 1. Assembly and sequencing of a comprehensive aptamer insertion library. a** Schematic showing the molecular cloning steps of saturated programmable insertion engineering (SPINE). Briefly, the splicing ribozyme was computationally divided into thirds (sections), and pooled oligos were synthesized for each section. Oligos contain the same sequence as their corresponding section, except that they have a genetic handle inserted at a specified site. The genetic handle consists of two outward-facing BsaI restriction sites with a short spacer between them. Each oligo is also flanked by two inward-facing BbsI restriction sites and a primer binding site that is distinct from that of the oligos corresponding to the other sections. One oligo is designed for each genetic handle insertion site in the section, such that across the entire oligo pool, there is a genetic handle between every nucleotide. The oligos for a given section are amplified from the pool by PCR, then cloned into the expression vector by BbsI Golden Gate to make the ‘staging library’. After transforming and extracting the staging library to amplify it, a second BsaI Golden gate is performed to replace the genetic handle with an aptamer. This process is repeated across all three sections to make a comprehensive library with aptamer insertions in every possible position in the ribozyme. **b** Abundance of variants containing different genetic handle insertions for each section in the staging library. **c** Abundance of variants containing different aptamer insertions across three replicates of the aptamer insertion library. Abundances were calculated using NGS of plasmid libraries, and the read counts were processed using custom scripts.

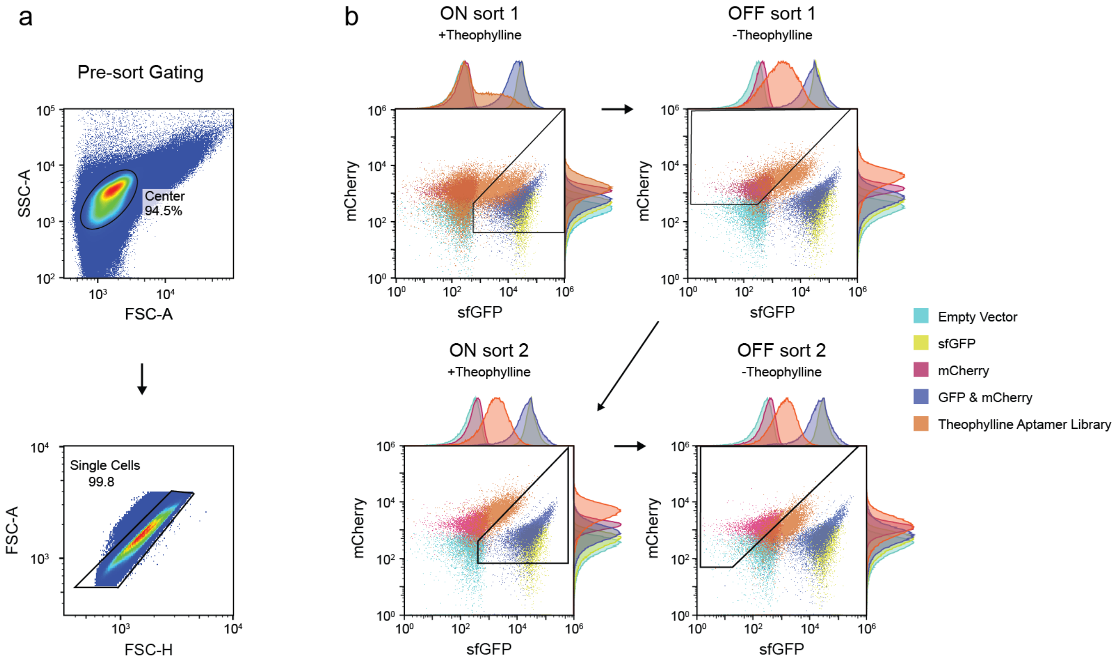

**Supplementary Fig. 2. Gating strategy for the theophylline aptamer domain insertion library. a** Representative gating strategy for FACS-seq pipeline. First, to obtain single cells, the densest region of the forward scatter area (FSC-A) vs side scatter area (SSC-A) was gated to eliminate outliers, cell clumps, and debris in the media. Then, doublet discrimination was performed by gating on forward scatter height (FSC-H) vs FSC-A. **b** Sorting gates for each round of library sorting for the theophylline aptamer. Libraries were sorted based on the sfGFP/mCherry expression ratio of the cells. High ratios were collected for ON sorts, low ratios for OFF sorts. For clarity, only one representative library of the three replicates is shown. Note that subsequent libraries only underwent the first two (fluoride) or three (ZMP) rounds of sorting.

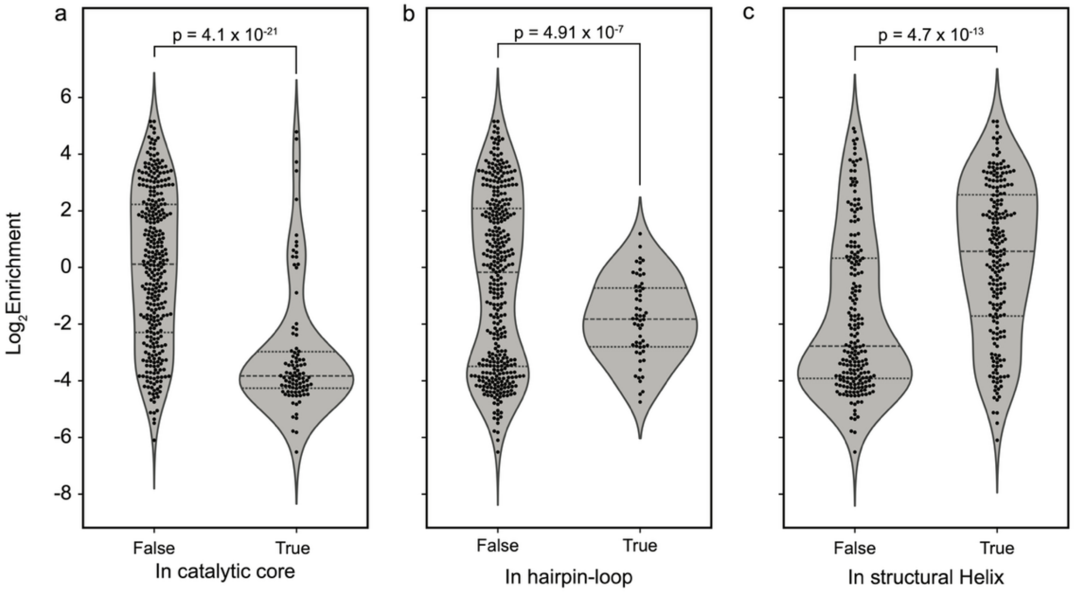

**Supplementary Fig. 3**. **The structural context of the insertion site of the theophylline aptamer within the splicing ribozyme influences enrichment outcomes**. Violin plots show the distribution of enrichment values for variants grouped by whether the aptamer was inserted within **a** the catalytic core, **b** a hairpin loop, or **c** a structural helix of the ribozyme. Individual enrichment values are shown as dots inside the violin plots. Heteroscedastic, two-tailed T-tests were performed on each pair to find the p values indicated above each pair.

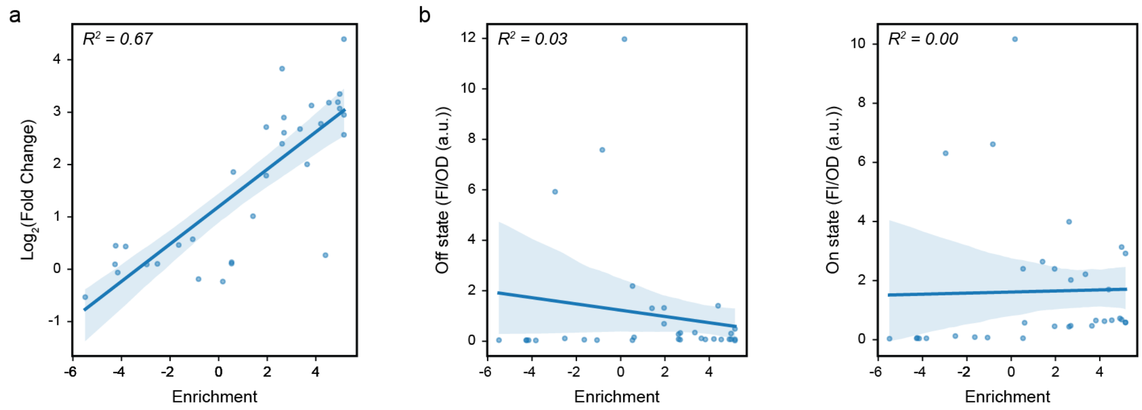

**Supplementary Fig. 4. Relationship between library enrichment and measured splicing activity of individual LASR variants.** Correlation between the enrichment and the fluorescence characterization data of individual variants in *E. coli* cells. Correlation between enrichment and (**a**) fold change (on/off), (**b**) off state in the absence of theophylline, and (**c**) on state in the presence of theophylline. For all plots, points represent the average of 4 biological replicates, and the shaded region indicates the 95% CI of the regression line.

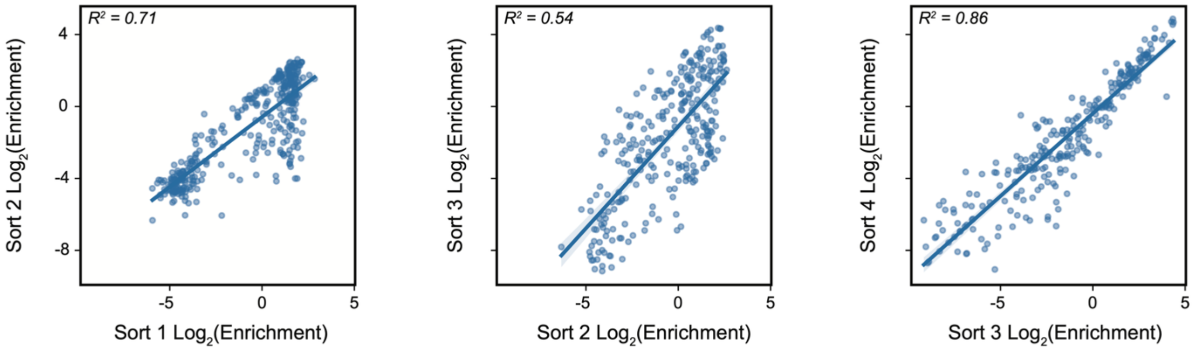

**Supplementary Fig. 5. Correlation of enrichment between rounds of different rounds of sorting.** The correlation of the enrichment of each insertion site relative to the starting library between each successive round of sorting for a single replicate of the theophylline aptamer library. Sorts 1 and 3 were ON sorts (*i.e.,* library was cultured in the presence of theophylline and variants with a high level GFP/mCherry expression ratio were selected). Sorts 2 and 4 were OFF sorts. As sorts 3 and 4 showed a strong correlation ($R^2$=0.84), this indicated minimal change in the enrichment scores with this additional sort, suggesting it was of little benefit. As such, reduced rounds of sorting were performed for the subsequent aptamer insertion libraries.

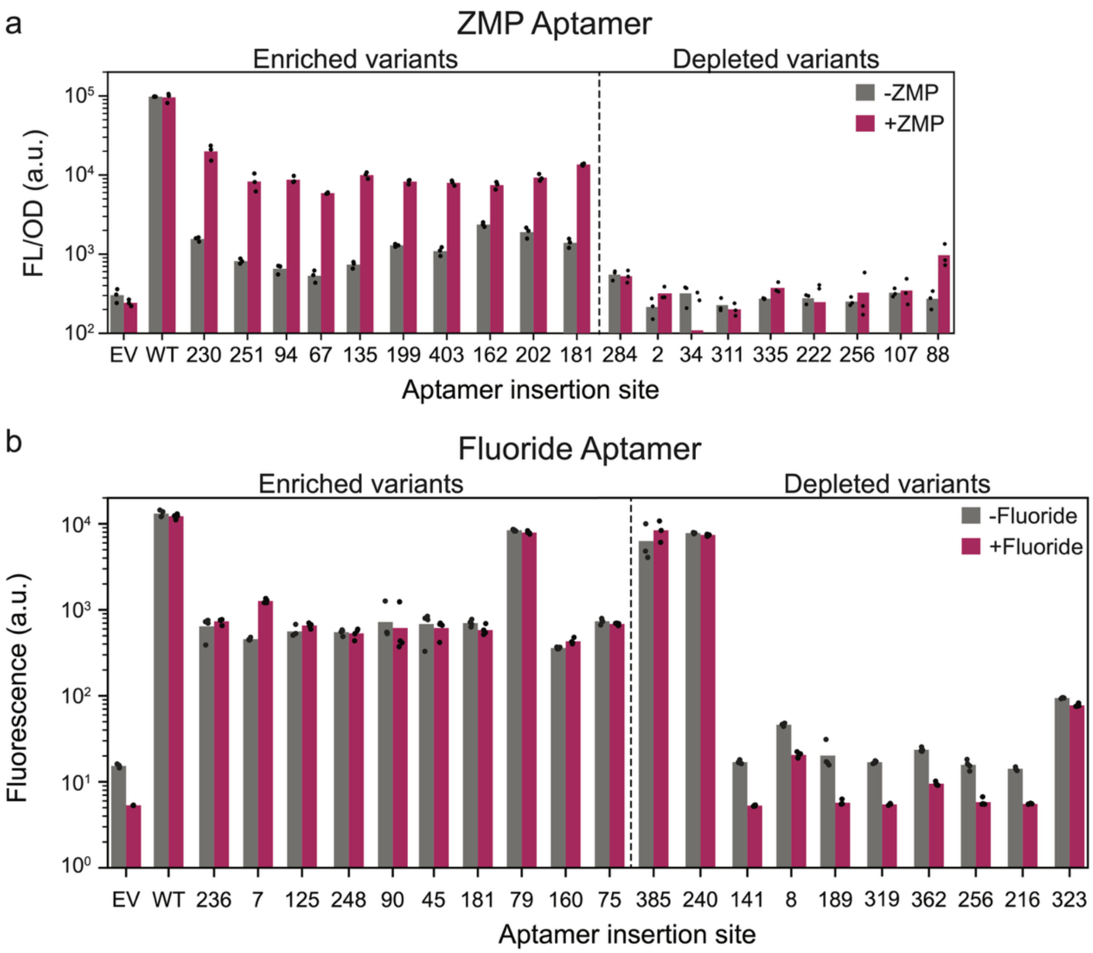

**Supplementary Figure 6. Individual characterization of enriched and depleted Fluoride and ZMP responsive LASRs**. **a** Splicing activity (as measured by GFP expression) of individually cloned enriched and depleted ZMP aptamer insertion variants. Data shows bulk fluorescence characterization (measured in units of fluorescence [FL]/optical density [OD] at 600 nm) in the presence and absence of 1 mM ZMP. **b** Splicing activity of enriched and depleted fluoride variants. Data shows single-cell fluorescence measured in the presence and absence of 200 μM fluoride. Bars indicate the mean of 3 biological replicates shown as points

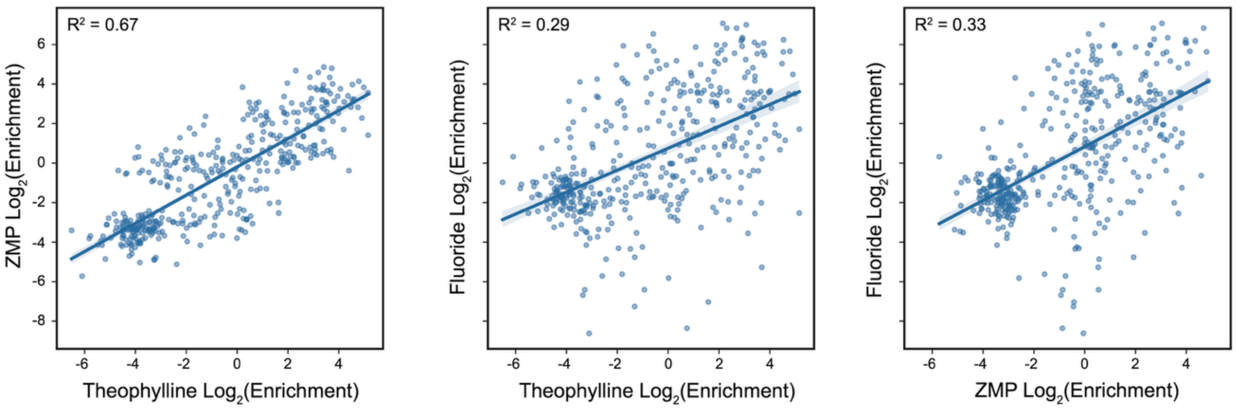

**Supplementary Fig. 7. Correlation of enrichment between the three aptamers.** The correlation of the mean enrichment of each insertion site between the aptamers binding theophylline, ZMP, and fluoride.

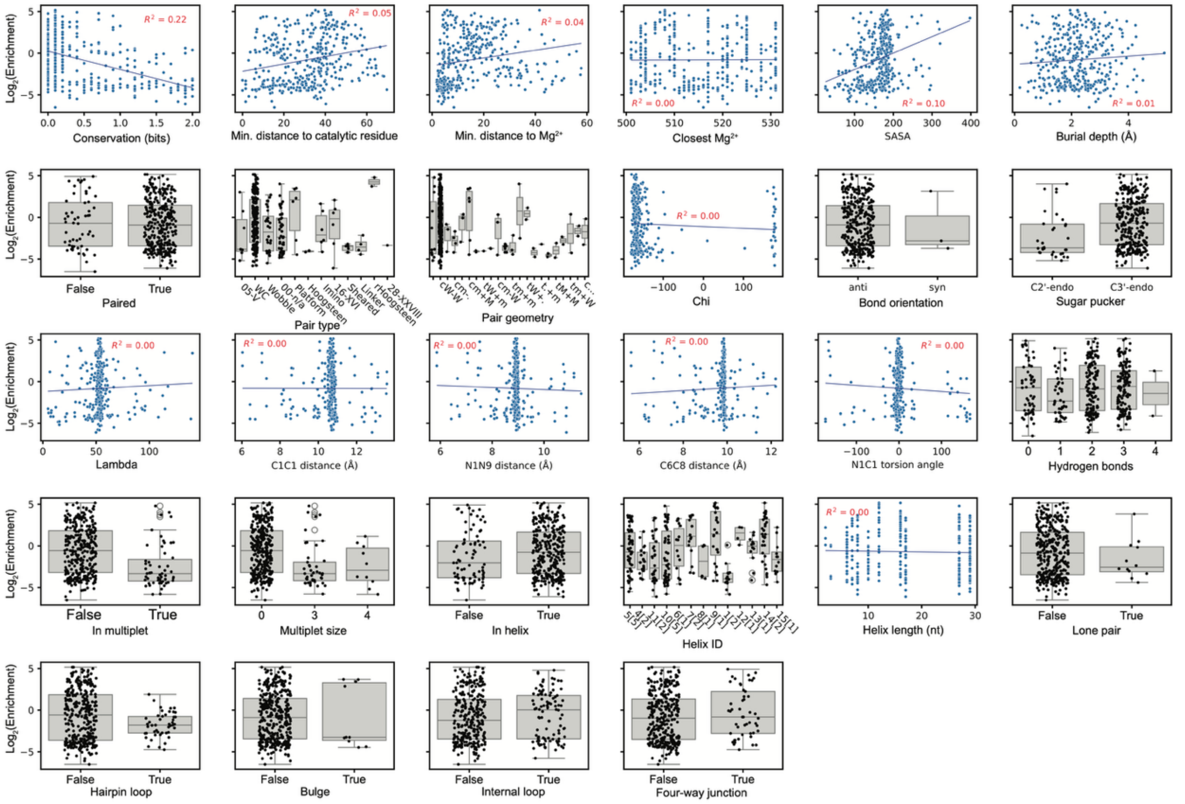

**Supplementary Fig. 8. Relationship between insert site enrichment and various structural parameters of the insert site.** Continuous parameters are shown as a scatterplot, discrete parameters are shown as a boxplot with strip plot overlay. SASA: solvent accessible surface area. Structural features are based on splicing ribozyme structure obtained by Su et al. Nature (2021)[1]. Conservation data based on the work of Che and Knight, NAR (2010)[2].

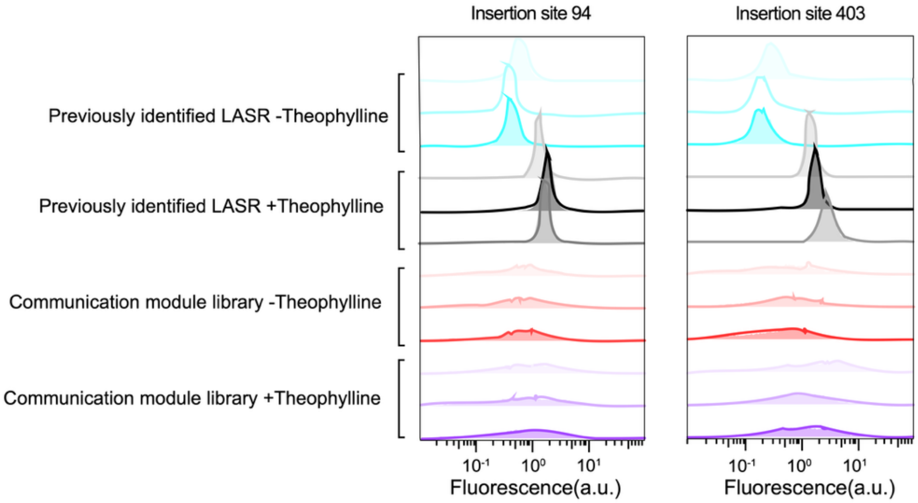

**Supplementary Fig. 9. Single-cell fluorescence distributions of the communication module library.** Fluorescence distributions of the communication module library for aptamer insertion site 94 and 403 in the presence and absence of theophylline are compared to the previously identified designed and show a broad range of activity levels. Data shows histograms of 3 biological replicates. The gating strategy used is described in Supplementary Figure 2.

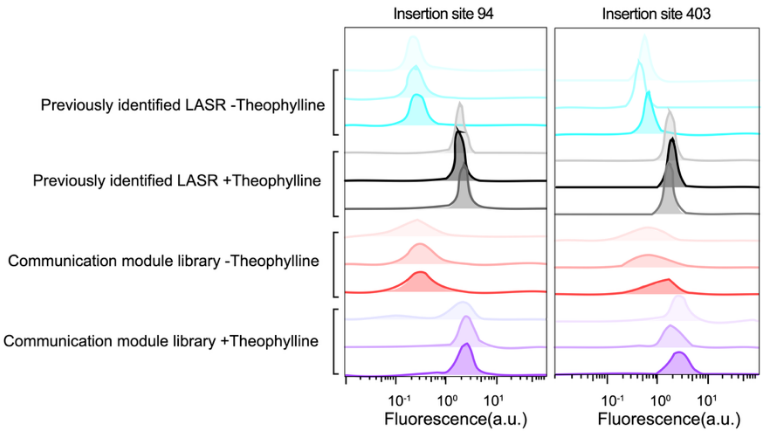

**Supplementary Fig. 10. Single-cell fluorescence distributions of the communication module library after cell sorting.** Fluorescence distributions of the communication module library after sorting for aptamer insertion site 94 and 403 in the presence and absence of theophylline are compared to the previously identified designed. Data shows histograms of 3 biological replicates. The gating strategy used is described in Supplementary Figure 2.

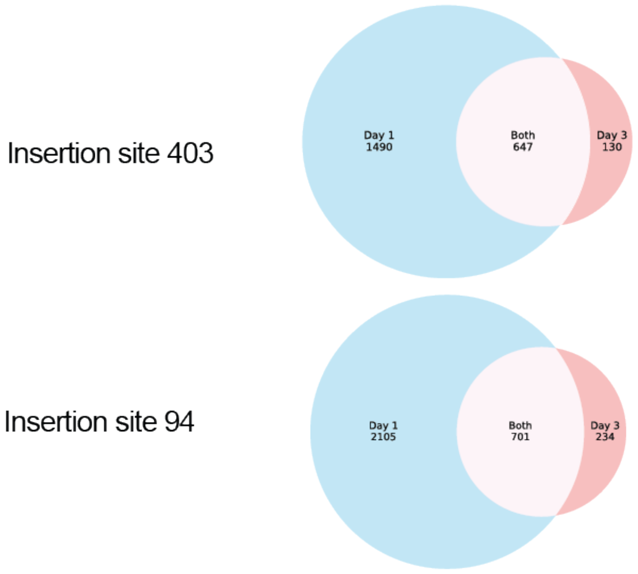

**Supplementary Fig. 11. Communication module diversity before and after sorting.** Venn diagrams comparing variant libraries pre-sorting (Day 1, blue) and post-sorting (Day 3, red) for aptamer insertion sites 403 and 94. The large overlap indicates that the majority of variants isolated on Day 3 (83% for site 403; 75% for site 94) were present in the initial Day 1 library, demonstrating successful enrichment of the starting population. Data shows the average number of variants across 3 biological replicates.

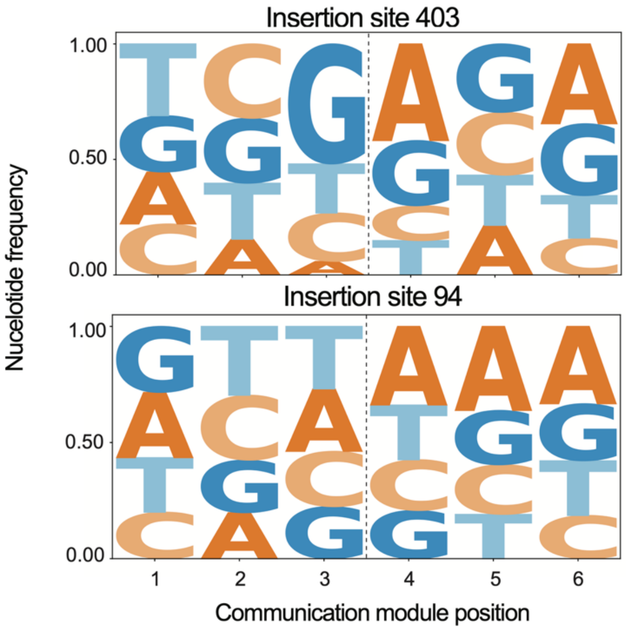

**Supplementary Fig. 12. Nucleotide frequency in the communication library for aptamer insertion sites 403 and 94 before sorting.** Sequence-based frequency logo plots of both aptamer insertion sites reveal that the naive library had even nucleotide distributions. Data shows the nucleotide frequency from 3 biological replicates.

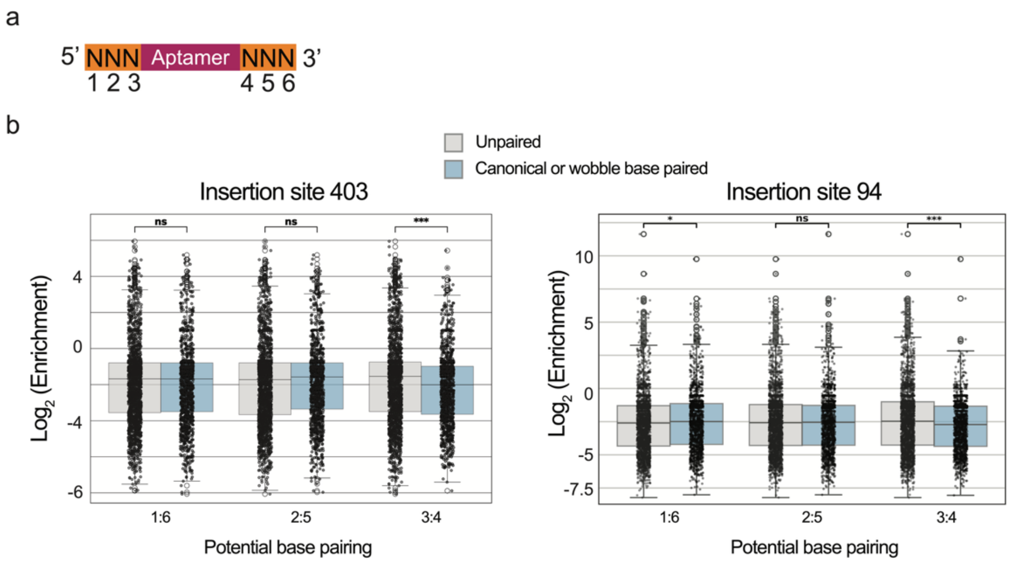

**Supplementary Fig. 13. Base-pairing trends across all three positions in the communication module libraries for aptamer insertion sites 403 and 94. a** Schematic of the 5' and 3' communication module positions 1 to 6. **b** Grouped boxplots overlaid with individual data points showing the log2 fold change of communication module variants, categorized by their pairing status at three specific interaction sites (1:6, 2:5, and 3:4). Variants at each position were classified as either capable of base pairing (comprising both canonical and wobble interactions) or unpaired. These data suggest that maintaining a paired interaction at the inner-most node (position 3:4) is unfavorable for enrichment in both aptamer insertion site 403 and 94. The box represents the interquartile range with the median indicated by the center line, while the overlaid points display the raw data distribution. Statistical significance between the unpaired and paired groups at each position was determined using a two-sided Mann-Whitney U test (*$p < 0.05$, **$p < 0.01$, ***$p < 0.001$, ns = not significant).

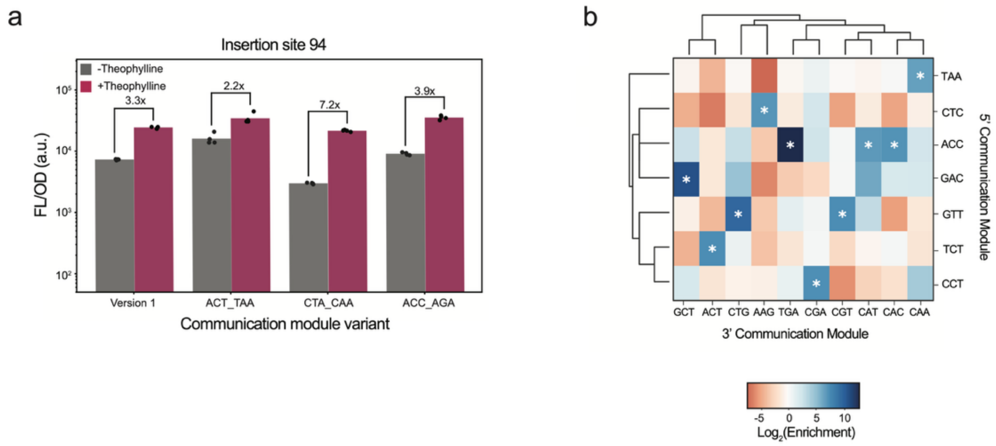

**Supplementary Fig. 14. Characterization of top communication modules and sequence-fitness landscape for insertion site 94. a** Characterization of the top three communication modules from insertion site 94 reveals robust ligand-switching activity. Bars indicate the mean of 4 biological replications shown as points. **b** Sequence-fitness landscapes of communication modules. The 5' and 3' communication modules from the top 10 enriched variants at insertion site 93 were identified (marked by asterisks), and a combinatorial matrix was constructed displaying corresponding enrichment scores.

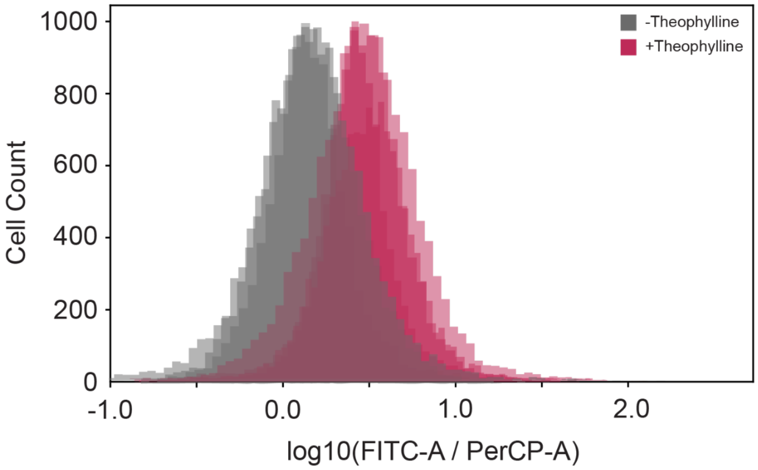

**Supplementary Fig. 15. Single-cell fluorescence characterization of LASR activity in *S. cerevisiae* using flow cytometry.** Splicing activity (as measured by GFP expression) of theophylline-responsive (LASR Theo-148) in *S. cerevisiae*. Data is single-cell fluorescence measured by flow cytometry in the presence and absence of 1 mM theophylline. GFP fluorescence was normalized to a constitutively expressed RFP. Data show a histogram of 4 biological replicates.

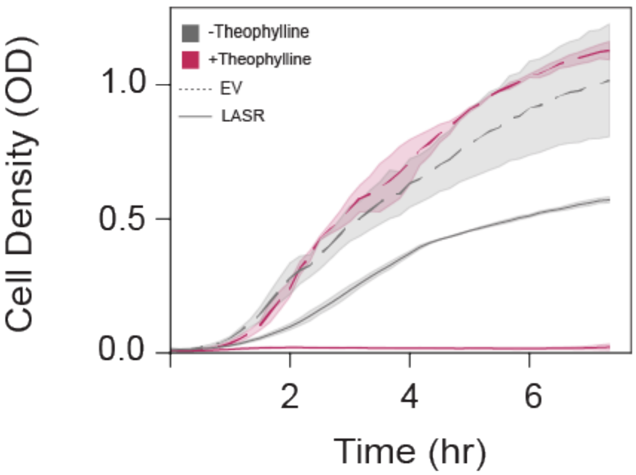

**Supplementary Fig. 16. Growth curve depicting regulation of a toxin gene, *ccdb* using the modular LASR design.** Cell growth is measured by optical density at 600 nm (OD600) for *E. coli* cells transformed with an empty vector control (EV) or a plasmid encoding LASR Theo-148 in the presence or absence of 1 mM theophylline. Data shows mean and standard deviation of 3 biological replicates.

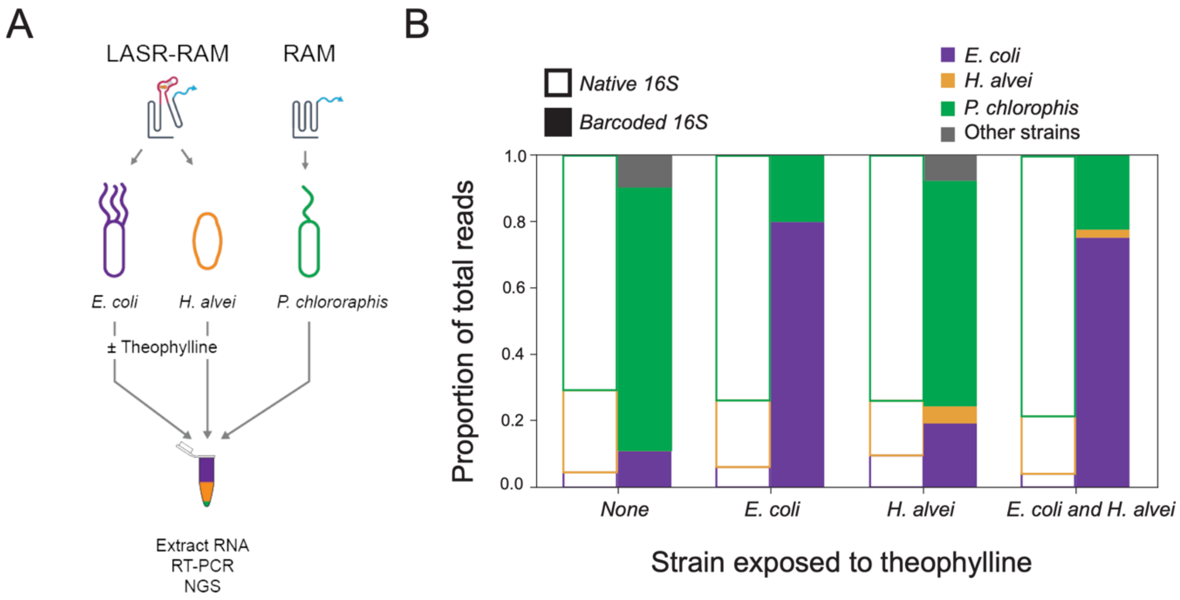

**Supplementary Fig. 17. Coupling LASR to an RNA recorder enables recording of intracellular chemical information in a synthetic microbial consortia. a** LASR-RAM system designed to splice synthetic RNA barcodes onto host 16S rRNA only when theophylline is present. In the consortia experiment, *E. coli* and *H. alvei* contain LASR-RAM and are cultured independently in the presence or absence of theophylline. They are then combined at an equal ratio (by optical density at 600 nm) immediately prior to RNA extraction. *P. chlororaphis* is transformed with a constitutively active RAM that barcodes 16S rRNA and added as a reference. Barcoded and native 16S rRNA are then reverse transcribed and sequenced by NGS. **b** Relative proportion of barcoded and native 16S rRNA sequencing reads for different consortia conditions where *E. coli* and/or *H. alvei* were exposed to theophylline. Data shows the mean of 3 biological replicates.

### Supplementary Table 1. Plasmids used in this study

| Plasmid Name | Description |
|---|---|
| pJEC1664 | Theophylline_148_CTT_GTT with pBBR1 origin |
| pJEC1635 | sfGFP with WT (constitutive) ribozyme inserted in the middle & constitutive mCherry |
| pJEC1636 | Constitutive mCherry |
| pJEC1637 | Theophylline LASR; insert site 403; 5' CM TTT; 3' CM CCC |
| pJEC1638 | Theophylline LASR; insert site 148; 5' CM CTT; 3' CM GTT |
| pJEC1639 | Theophylline LASR; insert site 163; 5' CM CTG; 3' TCC |
| pJEC1640 | Theophylline LASR; insert site 254; 5' CM AAT; 3' CM TGT |
| pJEC1641 | Theophylline LASR; insert site 182; 5' CM CCG; 3' CM ACC |
| pJEC1642 | Theophylline LASR; insert site 372; 5' CM CTG; 3' CM GAT |
| pJEC1643 | Theophylline LASR; insert site 413; 5' CM ACT; 3' CM TCC |
| pJEC1644 | Theophylline LASR; insert site 16; 5' CM CTG; 3' CM GTC |
| pJEC1645 | Theophylline LASR; insert site 24; 5' CM GCT; 3' CM CGA |
| pJEC1646 | Theophylline LASR; insert site 94; 5' CM GCT; 3' CM TCT |
| pJEC1647 | Theophylline_125_CCT_CCC |
| pJEC1648 | Theophylline_338_CTT_CGT |
| pJEC1649 | Theophylline_113_CTG_CCC |
| pJEC1650 | Theophylline_357_CGC_ACC |
| pJEC1651 | Theophylline_60_AGT_TCT |
| pJEC1652 | Theophylline_76_AAA_ACT |
| pJEC1653 | Theophylline_279_GTG_TCT |
| pJEC1654 | Theophylline LASR; insert site 38; 5' CM GTG; 3' CM CCC |
| pJEC1655 | Theophylline_163_CGT_GAT |
| pJEC1656 | Theophylline_403_TAG_GAA |
| pJEC1657 | Theophylline_375_CCG_TCC |
| pJEC1658 | Theophylline_24_GCT_TCT |
| pJEC1659 | Theophylline_160_CCG_TCC |
| pJEC1660 | Theophylline_160_CTT_CCC |
| pJEC1661 | Theophylline_362_CAA_CCC |
| pJEC1662 | Theophylline LASR; insert site 201; 5' CM CTG; 3' CM TAG |
| pJEC1663 | Theophylline LASR; insert site 275; 5' CM ACT; 3' CM ACC |
| pJEC1664 | Theophylline LASR; insert site 315; 5' CM CGT; 3' CM TAT |
| pJEC1665 | Theophylline LASR; insert site 80; 5' CM CTA; 3' CM CCG |
| pJEC1666 | Theophylline LASR; insert site 101; 5' CM CGT; 3' CM CCT |
| pJEC1667 | Theophylline LASR; insert site 107; 5' CM CAG; 3' CM CCC |
| pJEC1668 | Theophylline LASR; insert site 137; 5' CM CCT; 3' CM TAC |
| pJEC1669 | Theophylline LASR; insert site 414; 5' CM CCC; 3' CM TCT |

| | |
|---|---|
| pJEC1670 | ZMP_230_AAG_TTC |
| pJEC1671 | ZMP_251_CAT_GTG |
| pJEC1672 | ZMP_94_AAC_ATC |
| pJEC1673 | ZMP_67_ATT_GCC |
| pJEC1674 | ZMP_135_CAG_ATC |
| pJEC1675 | ZMP_199_GAC_ATC |
| pJEC1676 | ZMP_162_ATT_TTG |
| pJEC1677 | ZMP_403_GAG_CTC |
| pJEC1678 | ZMP_202_AAC_GTG |
| pJEC1679 | ZMP_181_GTG_GAG |
| pJEC1680 | ZMP_284_GAT_GGC |
| pJEC1681 | ZMP_110_CAC_GTC |
| pJEC1682 | ZMP_155_ACT_AGC |
| pJEC1683 | ZMP_371_AAA_GTG |
| pJEC1684 | ZMP_134_AAG_GAG |
| pJEC1685 | ZMP_222_TAT_GTG |
| pJEC1686 | ZMP_139_GGA_TAC |
| pJEC1687 | ZMP_406_GAA_TTC |
| pJEC1688 | ZMP_165_TAT_GAG |
| pJEC1689 | ZMP_182_AAA_AAG |
| pJEC1690 | ZMP_2_TAA_ATG |
| pJEC1691 | ZMP_34_TCG_TTG |
| pJEC1692 | ZMP_311_GCT_ACG |
| pJEC1693 | ZMP_335_GGC_ATC |
| pJEC1694 | ZMP_256_GAG_GTG |
| pJEC1695 | ZMP_107_CCC_GTC |
| pJEC1696 | ZMP_88_AGG_GTG |
| pJEC1697 | ZMP_403_AGG_GTG |
| pJEC1698 | Fluoride_236_TCC_AAT |
| pJEC1699 | Fluoride_7_TGG_ATC |
| pJEC1700 | Fluoride_125_GTC_CAT |
| pJEC1701 | Fluoride_248_CGC_GGC |
| pJEC1702 | Fluoride_90_GAC_GTT |
| pJEC1703 | Fluoride_79_TAA_CAT |
| pJEC1704 | Fluoride_181_ATA_TTT |
| pJEC1705 | Fluoride_160_ACC_GTT |
| pJEC1706 | Fluoride_45_ACG_AAC |
| pJEC1707 | Fluoride_75_AAG_GTC |
| pJEC1708 | Fluoride_385_TAC_GCT |

| | |
|---|---|
| pJEC1709 | Fluoride_240_GCA_TGT |
| pJEC1710 | Fluoride_141_GTC_TCC |
| pJEC1711 | Fluoride_8_ACC_TAT |
| pJEC1712 | Fluoride_189_AAC_CCG |
| pJEC1713 | Fluoride_319_GTG_GCC |
| pJEC1714 | Fluoride_362_TTC_GGC |
| pJEC1715 | Fluoride_256_TAC_CGC |
| pJEC1718 | pBBR1 theophylline LASR (broad host) |
| pJEC1754 | Fluoride_323_TCT_GCT |
| pJEC103 | SpecR empty vector (EV) |
| pJEC753 | sfGFP with WT (constitutive) ribozyme inserted in the middle |
| pJEC905 | pBBR1+KanR |
| pJEC906 | pBBR1+KanR+GFP |
| pJEC1755 | Theophylline_94_ACT_TAA |
| pJEC1756 | Theophylline_94_CTA_CAA |
| pJEC1757 | Theophylline_94_ACC_AGA |
| pJEC1758 | Theophylline_403_TCG_AGA |
| pJEC1759 | Theophylline_403_TCG_AAC |
| pJEC1760 | Theophylline_403_GTG_AAC |
| pJEC1761 | Theophylline_148_CTT_GTT with codon-optimized sfGFP and p10 loop for L. plantarum |
| pJEC1762 | Theophylline_148_CTT_GTT for yeast |
| pJEC1763 | Modular LASR Design 1 |
| pJEC1764 | Modular LASR Design 2 |
| pJEC1765 | Theophylline LASR with FMO output |
| pJEC1766 | Theophylline LASR with MHT output |
| pJEC1767 | Theophylline LASR with ccdB output |
| pJEC1654 | Inducible RAM plasmid |
| pJEC1244 | Constitutive RAM plasmid |
| pAOS191 | WT fluoride riboswitch controlling sfGFP |

### Supplementary Table 2. Example plasmid sequence

| | |
|---|---|
| Example of cis-splicing ribozyme with theophylline aptamer inserted at position 148, inserted within sfGFP [SpecR - CDF - PJ23119 - HP14 - RBS - sfGFP(1-66nt) - P1_loop - IGS - Ribozyme(1-148nt) - Theophylline aptamer- Ribozyme(148-387) sfGFP(66nt2-239) - TrrnB] | TTATTTGCCGACTACCTTGGTGATCTCGCCTTTCACGTAGTGGACAAATTCTTCCAA<br>CTGATCTGCGCGCGAGGCCAAGCGATCTTCTTCTTGTCCAAGATAAGCCTGTCTAG<br>CTTCAAGTATGACGGGCTGATACTGGGCCGGCAGGCGCTCCATTGCCCAGTCGG<br>CAGCGACATCCTTCGGCGCGATTTTGCCGGTTACTGCGCTGTACCAAATGCGGGA<br>CAACGTAAGCACTACATTTCGCTCATCGCCAGCCCAGTCGGGCGGCGAGTTCCAT<br>AGCGTTAAGGTTTCATTTAGCGCCTCAAATAGATCCTGTTCAGGAACCGGATCAAA<br>GAGTTCCTCCGCCGCTGGACCTACCAAGGCAACGCTATGTTCTCTTGCTTTTGTCA<br>GCAAGATAGCCAGATCAATGTCGATCGTGGCTGGCTCGAAGATACCTGCAAGAAT<br>GTCATTGCGCTGCCATTCTCCAAATTGCAGTTCGCGCTTAGCTGGATAACGCCACG<br>GAATGATGTCGTCGTGCACAACAATGGTGACTTCTACAGCGCGGAGAATCTCGCT<br>CTCTCCAGGGGAAGCCGAAGTTTCCAAAAGGTCGTTGATCAAAGCTCGCCGCGTT<br>GTTTCATCAAGCCTTACGGTCACCGTAACCAGCAAATCAATATCACTGTGTGGCTT<br>CAGGCCGCCATCCACTGCGGAGCCGTACAAATGTACGGCCAGCAACGTCGGTTC<br>GAGATGGCGCTCGATGACGCCAACTACCTCTGATAGTTGAGTCGATACTTCGGCG<br>ATCACCGCTTCCCTCATACTCTTCCTTTTTCAATATTATTGAAGCATTTATCAGGGTT<br>ATTGTCTCATGAGCGGATACATATTTGAATGTATTTAGAAAAATAAACAAATAGCTA<br>GCTCACTCGGTCGCTACGCTCCGGGCGTGAGACTGCGGCGGGCGCTGCGGACAC<br>ATACAAAGTTACCCACAGATTCCGTGGATAAGCAGGGGACTAACATGTGAGGCAA<br>AACAGCAGGGCCGCGCCGGTGGCGTTTTTCCATAGGCTCCGCCCTCCTGCCAGA<br>GTTCACATAAACAGACGCTTTTCCGGTGCATCTGTGGGAGCCGTGAGGCTCAACC<br>ATGAATCTGACAGTACGGGCGAAACCCGACAGGACTTAAAGATCCCCACCGTTTC<br>CGGCGGGTCGCTCCCTCTTGCGCTCTCCTGTTCCGACCCTGCCGTTTACCGGATA<br>CCTGTTCCGCCTTTCTCCCTTACGGGAAGTGTGGCGCTTTCTCATAGCTCACACAC<br>TGGTATCTCGGCTCGGTGTAGGTCGTTCGCTCCAAGCTGGGCTGTAAGCAAGAAC<br>TCCCCGTTCAGCCCGACTGCTGCGCCTTATCCGGTAACTGTTCACTTGAGTCCAAC<br>CCGGAAAAGCACGGTAAAACGCCACTGGCAGCAGCCATTGGTAACTGGGAGTTCG<br>CAGAGGATTTGTTTAGCTAAACACGCGGTTGCTCTTGAAGTGTGCGCCAAAGTCCG<br>GCTACACTGGAAGGACAGATTTGGTTGCTGTGCTCTGCGAAAGCCAGTTACCACG<br>GTTAAGCAGTTCCCCAACTGACTTAACCTTCGATCAAACCACCTCCCCAGGTGGTT<br>TTTTCGTTTACAGGGCAAAAGATTACGCGCAGAAAAAAAGGATCTCAAGAAGATCC<br>TTTGATCTTTTCTACTGAACCGCTCTAGATTTCAGTGCAATTTATCTCTTCAAATGTA<br>GCACCTGAAGTCAGCCCCATACGATATAAGTTGTAATTCTCATGTTAGTCATGCCC<br>CGCGCCCACCGGAAGGAGCTGACTGGGTTGAAGGCTCTCAAGGGCATCGGTCGA<br>GATCCCGGTGCCTAATGAGTGAGCTAACTTACACCCGCAGTTCCAGTACGGTTCC<br>ATTAATTGCGTTGCGCTGCTTCCGGCTGAATTCTAAAGATCTTTGACAGCTAGCTC<br>AGTCCTAGGTATAATACTAGTACGTCGACTCTCGAGTGAGATTGTTGACGGTACCG<br>TATTTTGGATCTAGGAGGAAGGATCTATGAGCAAAGGAGAAGAACTTTTCACTGGA<br>GTTGTCCCAATTCTTGTTGAATTAGATGGTGATGTTAATGGGCACAAATTTTCTGTC<br>CGTGGAGAGGGTGAAGGTGATGCTACAAACGGAAAACTCACCCTTAAATTTATTTG<br>CACTACTGGAAAACTACCTGTTCCGTGGCCAACACTTGTCACTACTCTGACCTAAA<br>TAGCAATATTTACCTTTGGGTCATCAGGCATGCACCTGGTAGCTAGTCTTTAAACC<br>AATAGATTGCATCGGTTTAAAAGGCAAGACCGTCAAATTGCGGGAAAGGGGTCAA<br>CAGCCGTTCAGTACCAAGTCAGCGCTTGGTGATACCAGCATCGTCTTGATGCCCT<br>TGGCAGCACCGTTGAGCTCAGGGGAAACTTTGAGATGGCCTTGCAAAGGGTATGG<br>TAATAAGCTGACGGACATGGTCCTAACCACGCAGCCAAGTCCTAAGTCAACAGATC<br>TTCTGTTGATATGGATGCAGTTCACAGACTAAATGTCGGTCGGGGAAGATGTATTCT<br>TCTCATAAGATATAGTCGGACCTCTCCTTAATGGGAGCTAGCGGATGAAGTGATGCA<br>ACACTGGAGCCGCTGGGAACTAATTTGTATGCGAAAGTATATTGATTAGTTTTGGAGT<br>ACTCGAAACGGCATGACTTTTTCAAGAGTGCCATGCCCGAAGGTTATGTACAGGAACG<br>CACTATATCTTTCAAAGATGACGGGACCTACAAGACGCGTGCTGAAGTCAAGTTTGAA<br>GGTGATACCCTTGTTAATCGTATCGAGTTAAAGGGTATTGATTTTAAAGAAGATGGA<br>AACATTCTTGGACACAAACTCGAGTACAACTTTAACTCACACAATGTATACATCACG<br>GCAGACAAACAAAAGAATGGAATCAAAGCTAACTTCAAAATTCGCCACAACGTTGA<br>AGATGGTTCCGTTCAACTAGCAGACCATTATCAACAAAATACTCCAATTGGCGATG<br>GCCCTGTCCTTTTACCAGACAACCATTACCTGTCGACACAATCTGTCCTTTCGAAA<br>GATCCCAACGAAAAGCGTGACCACATGGTCCTTCTTGAGTTTGTAACTGCTGCTGG<br>GATTACACATGGCATGGATGAGCTCTACAAATAAGGATCTCAAAGCCCGCCGAAAG<br>GCGGGCTTTTTTTT |

### Supplementary Table 3. Aptamer sequences used.

| | |
|---|---|
| Theophylline | GGTGATACCAGCATCGTCTTGATGCCCTTGGCAGCACC |
| ZMP | ATTAGTCATATGACTGACGGAAGTGGAGTTACCACATGAAGTATGACTAGGCATATTATCT TATATGCCACAAAAAGCCGACCGTCTGGGC |
| Fluoride | TTATAGGCGATGGAGTTCGCCATAAACGCTGCTTAGCTAATGACTCCTACCAG |

### Supplementary Table 4. Strains used.

| Species | Genotype / Strain |
|---|---|
| *Escherichia coli* | MG1655 |
| *Hafnia alvei* | HAMBI 1279 (ATCC 13337 T) |
| *Kluyvera intermedia* | HAMBI 1299 (ATCC 33110 T) |
| *Pseudomonas chlororaphis* | HAMBI 1977 (ATCC 17415) |
| *Comamonas testosteroni* | HAMBI 403 (ATCC 11996 T) |
| *Lactiplantibacillus plantarum* | WCFS1 |
| *Saccharomyces cerevisae* | BY4741 |
| *Escherichia coli* | ECAH5 |

### Supplementary Table 5. Primers used.

| Purpose | Primer Name | Sequence (5' to 3') |
| --- | --- | --- |
| For RT-qPCR | qRM05 | CTAGCTGGTCTGAGAGGATG |
| | qRM16 | TGTGCAATATTCCCCACTGC |
| | qRM14 | TGTAGGTCCCGTCATCTTTG |
| | qRM17 | AGTCGGAATCGCTAGTAATCG |
| | qPCR probe | CGGTGAATATGGTGTTCAATGCTTTTCCC |
| LASR-RAM RT primer | AOS17B | TACATAACCTTCGGGCATGG |
| | AOS56B | GACGGGCGGTGWGTRCA |
| | DSZ46F (Blocking Oligo) | CCCAGCGGCTCCAGTGTTGC |
| LASR-RAM PCR primer native 16S rRNA | 968F | AACGCGAAGAACCTTAC |
| | AOS56B | GACGGGCGGTGWGTRCA |
| LASR-RAM PCR primer barcoded 16S rRNA | 968F | AACGCGAAGAACCTTAC |
| | DSZ39E | TAACGGGAAAAGCATTGAAC |
| Adapter PCR primer native 16S rRNA | DSZ29C | acactctttccctacacgacgctcttccgatctgtttAACGCGAAGAACCTTAC |
| | DSZ31C-II | acactctttccctacacgacgctcttccgatctcaatAACGCGAAGAACCTTAC |
| | DSZ03E | gactggagttcagacgtgtgctcttccgatctgagcGACGGGCGGTGWGTRCA |
| | DSZ04E-II | gactggagttcagacgtgtgctcttccgatctcgttGACGGGCGGTGWGTRCA |
| Adapter PCR primer barcoded 16S rRNA | DSZ29C | acactctttccctacacgacgctcttccgatctgtttAACGCGAAGAACCTTAC |
| | DSZ30C | acactctttccctacacgacgctcttccgatctaacgAACGCGAAGAACCTTAC |
| | DSZ66C | gactggagttcagacgtgtgctcttccgatctacctGTCTGGGAAAAGCATTGAA C |
| | DSZ67C | gactggagttcagacgtgtgctcttccgatctgaaaGTCTGGGAAAAGCATTGAA C |
| SPINE cloning – backbone amplification | AOS03B | ATACGTCTCCAGAGTAGTGACAAGTGTTGGCc |
| | AOS04B | ATACGTCTCTACCAAGTCTCAGGGGAAACTTT |
| | AOS05B | ATACGTCTCGTACTGAACGGCTGTTGACCC |
| | AOS06B | ATACGTCTCCGGTCGGGGAAGATGTATTC |
| | AOS07B | ATACGTCTCCCGACATTTAGTCTGTGAACTGCA |
| | AOS08B | ATACGTCTCCGATGGTGTTCAATGCTTTTCCc |
| SPINE Cloning – oligo subpool amplification | AOS16B | ATATAGATGCCGTCCTAGCG |
| | AOS78A | TTTCCTGTGCCCATGGTAGTA |
| | AOS80A | TTGGTCATGTGCTTTTCGTTGG |
| | AOS81A | TCGGATACTCAGAATCGATTCC |
| | AOS01B | TCCGACGGGGAGTATATACTC |
| | AOS02B | TGCTTCAGGCCAAAATAATCTCC |
| iPCR primers to clone pAOS106 | AOS05D | TGATGCCCTTGGCAGCACCNNNGAGCAGAGACCatgcgagagtagg gaac |

| | | |
|---|---|---|
| (Theophylline aptamer part plasmid) | AOS06D | AGACGATGCTGGTATCACCNNNCGCTAGAGACCcaacttatatcgtatggggctg |
| iPCR primers to clone pAOS149 (ZMP aptamer part plasmid) | AOS54E | ACTAGGCATATTATCTTATATGCCACAAAAAGCCGACCGTCTGGGCNNNGAGCAGAGAccaggcatcaaataaaacg |
| | AOS55E | CATACTTCATGTGGTAACTCCACTTCCGTCAGTCATATGACTAATNNNCGCTAGAGACCcaacttatatcgtatggggc |
| SPINE Staging library PCR for amplicon NGS sequencing | AOS42B | ACACTCTTTCCCTACACGACGCTCTTCCGATCTccaacacttgtcactactc |
| | AOS43B | GACTGGAGTTCAGACGTGTGCTCTTCCGATCTggaaaagcattgaacaccat |
| SPINE aptamer library PCR for amplicon NGS sequencing - first half | AOS42B | ACACTCTTTCCCTACACGACGCTCTTCCGATCTccaacacttgtcactactc |
| | AOS81E | GACTGGAGTTCAGACGTGTGCTCTTCCGATCTttgacttaggacttggctg |
| SPINE aptamer library PCR for amplicon NGS sequencing - second half | AOS01F | ACACTCTTTCCCTACACGACGCTCTTCCGATCTtaataagctgacggacatgg |
| | AOS43B | GACTGGAGTTCAGACGTGTGCTCTTCCGATCTggaaaagcattgaacaccat |
| Cloning individual theophylline LASR variants (iPCR) | AOS23D | TGATGCCCTTGGCAGCACCCCCGAGCAGTTTTGGAGTACTCGatg |
| | AOS24D | AGACGATGCTGGTATCACCAAACGCTAATCAATATACTTTCGCATAC |
| | AOS25D | TGATGCCCTTGGCAGCACCGTTGAGCTCAGGGGAAACTTTGAGATGG |
| | AOS26D | AGACGATGCTGGTATCACCAAGCGCTGACTTGGTACTGAACGGCTG |
| | AOS27D | TGATGCCCTTGGCAGCACCTCCGAGCAGATGGCCTTGCAAAGGGTA |
| | AOS28D | AGACGATGCTGGTATCACCCAGCGCTCAAAGTTTCCCCTGAGACTT |
| | AOS29D | TGATGCCCTTGGCAGCACCTGTGAGCGGATGCAGTTCACAGACTAA |
| | AOS30D | AGACGATGCTGGTATCACCATTCGCTATATCAACAGAAGATCTGTTGAC |
| | AOS31D | TGATGCCCTTGGCAGCACCACCGAGCATGGTAATAAGCTGACGGAC |
| | AOS32D | AGACGATGCTGGTATCACCCGGCGCTACCCTTTGCAAGGCCATC |
| | AOS33D | TGATGCCCTTGGCAGCACCGATGAGCGAACTAATTTGTATGCGAAAgtatattgattagttttggag |
| | AOS34D | AGACGATGCTGGTATCACCCAGCGCTCCAGCGGCTCCAGTGTTG |
| | AOS35D | TGATGCCCTTGGCAGCACCTCCGAGCTACTCGATGGTGTTCAATGC |
| | AOS36D | AGACGATGCTGGTATCACCAGTCGCTCTCCAAAACTAATCAATATActttcgcatac |
| | AOS37D | TGATGCCCTTGGCAGCACCGTCGAGCATTTACCTTTGGGTCAAAAA |
| | AOS38D | AGACGATGCTGGTATCACCCAGCGCTATTGCTATTTAGGTCAGAGT |
| | AOS39D | TGATGCCCTTGGCAGCACCCGAGAGCTTGGGTCAAAAAGTTATCAG |
| | AOS40D | AGACGATGCTGGTATCACCAGCCGCTAGGTAAATATTGCTATTTAGgtc |
| | AOS41D | TGATGCCCTTGGCAGCACCTCTGAGCAGGCAAGACCGTCAAATTGC |
| | AOS42D | AGACGATGCTGGTATCACCAGCCGCTTTTAAACCGATGCAATCTATtggtttaaag |
| | AOS22E | TGATGCCCTTGGCAGCACCCCCGAGCCAACAGCCGTTCAGTACC |

| | | |
|---|---|---|
| | AOS23E | AGACGATGCTGGTATCACCCGCTACCCCTTTCCCGCAA |
| | AOS24E | TGATGCCCTTGGCAGCACCGAGCCGGGAAAGGGGTCAACA |
| | AOS25E | AGACGATGCTGGTATCACCCAGCGCTCAATTTGACGGTCTTGCCTTT |
| | AOS26E | TGATGCCCTTGGCAGCACCTCTGAGCTAGTCTTTAAACCAATAGATTGCATC |
| | AOS27E | AGACGATGCTGGTATCACCACTCGCTGCTACCAGGTGCATGC |
| | AOS28E | TGATGCCCTTGGCAGCACCTCTGAGCGGTCGGGGAAGATGTATTCTT |
| | AOS29E | AGACGATGCTGGTATCACCCACCGCTGACATTTAGTCTGTGAACTGCATC |
| | AOS30E | TGATGCCCTTGGCAGCACCTCTGAGCTTGGGTCAAAAAGTTATCAGGCAT |
| | AOS31E | AGACGATGCTGGTATCACCAGCCGCTAGGTAAATATTGCTATTTAGGTCAGAGTAGT |
| | AOS32E | TGATGCCCTTGGCAGCACCCCCGAGCTTGAGATGGCCTTGCAAAG |
| | AOS33E | AGACGATGCTGGTATCACCAAGCGCTAGTTTCCCCTGAGACTTGG |
| | AOS34E | TGATGCCCTTGGCAGCACCCCCGAGCGAGCCGCTGGGAACTA |
| | AOS35E | AGACGATGCTGGTATCACCTTGCGCTCAGTGTTGCATCACTTCATCC |
| | AOS38E | TGATGCCCTTGGCAGCACCTAGGAGCCATGGTCCTAACCACGC |
| | AOS39E | AGACGATGCTGGTATCACCCAGCGCTTCCGTCAGCTTATTACCATACC |
| | AOS40E | TGATGCCCTTGGCAGCACCACCGAGCTGTCGGTCGGGGAAG |
| | AOS41E | AGACGATGCTGGTATCACCAGTCGCTTTTAGTCTGTGAACTGCATCCA |
| | AOS42E | TGATGCCCTTGGCAGCACCTATGAGCCGGACCTCTCCTTAATGGG |
| | AOS43E | AGACGATGCTGGTATCACCACGCGCTACTATATCTTATGAGAAGAATACATCTTCCCC |
| | AOS44E | TGATGCCCTTGGCAGCACCCCGGAGCTGCATCGGTTTAAAAGGCAAG |
| | AOS45E | AGACGATGCTGGTATCACCTAGCGCTATCTATTGGTTTAAAGACTAGCTACCAGG |
| | AOS46E | TGATGCCCTTGGCAGCACCCCTGAGCACCGTCAAATTGCGGGAA |
| | AOS47E | AGACGATGCTGGTATCACCACGCGCTCTTGCCTTTTAAACCGATGCAAT |
| | AOS48E | TGATGCCCTTGGCAGCACCCCCGAGCAAATTGCGGGAAAGGGG |
| | AOS49E | AGACGATGCTGGTATCACCCTGCGCTGACGGTCTTGCCTTTTAAACC |
| | AOS52E | TGATGCCCTTGGCAGCACCTCTGAGCACTCGATGGTGTTCAATGCTT |
| | AOS53E | AGACGATGCTGGTATCACCGGGCGCTACTCCAAAACTAATCAATATACTTTCGCA |
| Cloning individual theophylline LASR variants (Golden Gate) | AOS47D | GATACCAGCATCGTCTTGATGCCCTTGG |
| | AOS48D | GCTGCCAAGGGCATCAAGACGATGCTGG |
| | AOS53D | GCCAGGTCTCCCAGCACCCGTGAGCAGCGGATGAAGTGATGCAACA |
| | AOS54D | GCCAGGTCTCCTATCACCAAGCGCTAGCTCCCATTAAGGAGAGGTC |
| | AOS57D | TATAGGTCTCACAGCACCACCGAGCCACTGGAGCCGCTGG |

| | | |
|---|---|---|
| | AOS58D | GCCAGGTCTCCTATCACCGCGCGCTTTGCATCACTTCATCCGCTAG |
| | AOS61D | GCCAGGTCTCCCAGCACCACTGAGCAGATTGCATCGGTTTAAAAGGCAAGA |
| | AOS62D | GCCAGGTCTCCTATCACCTTTCGCTATTGGTTTAAAGACTAGCTACCAGGTG |
| | AOS65D | GATAGGTCTCCCAGCACCCCCGAGCTATCAGGCATGCACCTGGTAG |
| | AOS66D | GCCAGGTCTCCTATCACCCACCGCTACTTTTTGACCCAAAGGTAAATATTGCTATTTAGG |
| | AOS67D | GCTAGGTCTCCCAGCACCGATGAGCAGATGGCCTTGCAAAGGGTAT |
| | AOS68D | GCCAGGTCTCCTATCACCACGCGCTCAAAGTTTCCCCTGAGACTTGGTAC |
| | AOS69D | GCCAGGTCTCCCAGCACCGAAGAGCAGTTTTGGAGTACTCGATGGTGTTC |
| | AOS70D | GCCAGGTCTCCTATCACCCTACGCTAATCAATATACTTTCGCATACAAATTAGTTCCCAG |
| | AOS71D | GCCAGGTCTCCCAGCACCTCCGAGCCTAATTTGTATGCGAAAGTATATTGATTAGTTTTGGAGTAC |
| | AOS72D | GATAGGTCTCCTATCACCCGGCGCTTTCCCAGCGGCTCCA |
| | AOS75D | GATAGGTCTCCCAGCACCTCCGAGCTTGAGATGGCCTTGCAAAGG |
| | AOS76D | GCCAGGTCTCCTATCACCCGGCGCTAGTTTCCCCTGAGACTTGGTACT |
| | AOS16E | GCTAGGTCTCCCAGCACCTACGAGCAGTACCAAGTCTCAGGGGAAACT |
| | AOS17E | GCTAGGTCTCCTATCACCAGGCGCTGAACGGCTGTTGACCCC |
| Cloning pJEC1718 (broad host range theophylline LASR) | AOS06F | gagcctttcgttttatttgattatttgtagagctcatccatgc |
| | AOS07F | ggcagtgagcgcaacgcaatttgacagctagctcagtcct |
| | AOS08F | aggactgagctagctgtcaaattgcgttgcgctcactgcc |
| | AOS09F | tggatgagctctacaaataatcaaataaaacgaaaggctcagtcg |
| Cloning pAOS191 (WT fluoride riboswitch) | AOS41F | GCTAAGCAGCGTTTATGGCGAACTCCATCGCCtataaactagtattatacctaggactgagctagctg |
| | AOS42F | TAATGACTCCTACCAGTATCACTACTGGTAGGAGTCTATTTTTTTaggaggaaggatctatgagcaaag |
| Cloning individual ZMP LASR variants (iPCR) | AOS19G | CTAGGCATATTATCTTATATGCCACAAAAAGCCGACCGTCTGGGCTTCGAGCAGTCAACAGATCTTCTGTTGATATG |
| | AOS20G | TCATACTTCATGTGGTAACTCCACTTCCGTCAGTCATATGACTAATCTTCGCTTAGGACTTGGCTGCGTG |
| | AOS23G | CTAGGCATATTATCTTATATGCCACAAAAAGCCGACCGTCTGGGCGTGGAGCTATGGATGCAGTTCACAGAC |
| | AOS24G | TCATACTTCATGTGGTAACTCCACTTCCGTCAGTCATATGACTAATATGCGCTTCAACAGAAGATCTGTTGAC |
| | AOS68F | CTAGGCATATTATCTTATATGCCACAAAAAGCCGACCGTCTGGGCATCGAGCAGGCAAGACCGTCAAATTGC |
| | AOS69F | TCATACTTCATGTGGTAACTCCACTTCCGTCAGTCATATGACTAATGTTCGCTTTTAAACCGATGCAATCTATTG |
| | AOS62F | CTAGGCATATTATCTTATATGCCACAAAAAGCCGACCGTCTGGGCGCCGAGCTAAACCAATAGATTGCATCGG |
| | AOS63F | TCATACTTCATGTGGTAACTCCACTTCCGTCAGTCATATGACTAATAATCGCTAAGACTAGCTACCAGGTGCATG |
| | AOS76F | CTAGGCATATTATCTTATATGCCACAAAAAGCCGACCGTCTGGGCATCGAGCTCAGTACCAAGTCTCAGGGGAAAC |
| | AOS77F | TCATACTTCATGTGGTAACTCCACTTCCGTCAGTCATATGACTAATCTGCGCTACGGCTGTTGACCCCTTTC |
| | AOS11G | CTAGGCATATTATCTTATATGCCACAAAAAGCCGACCGTCTGGGCATCGAGCGACATGGTCCTAACCACGC |

| | | |
|---|---|---|
| | AOS12G | TCATACTTCATGTGGTAACTCCACTTCCGTCAGTCATATGACTAATGTCCGCTCGTCAGCTTATTACCATACCC |
| | AOS03G | CTAGGCATATTATCTTATATGCCACAAAAAGCCGACCGTCTGGGCTTGGAGCGAGATGGCCTTGCAAAGGG |
| | AOS04G | TCATACTTCATGTGGTAACTCCACTTCCGTCAGTCATATGACTAATAATCGCTAAAGTTTCCCCTGAGACTTGGTAC |
| | AOS41G | CTAGGCATATTATCTTATATGCCACAAAAAGCCGACCGTCTGGGCCTCGAGCAGTTTTGGAGTACTCGATGG |
| | AOS42G | TCATACTTCATGTGGTAACTCCACTTCCGTCAGTCATATGACTAATCTCCGCTAATCAATATACTTTCGCATAC |
| | AOS13G | CTAGGCATATTATCTTATATGCCACAAAAAGCCGACCGTCTGGGCGTGGAGCATGGTCCTAACCACGCAGC |
| | AOS14G | TCATACTTCATGTGGTAACTCCACTTCCGTCAGTCATATGACTAATGTTCGCTGTCCGTCAGCTTATTACCATACC |
| | AOS07G | CTAGGCATATTATCTTATATGCCACAAAAAGCCGACCGTCTGGGCGAGGAGCTATGGTAATAAGCTGACGGACATG |
| | AOS08G | TCATACTTCATGTGGTAACTCCACTTCCGTCAGTCATATGACTAATCACCGCTCCCTTTGCAAGGCCATCTC |
| | AOS29G | CTAGGCATATTATCTTATATGCCACAAAAAGCCGACCGTCTGGGCGGCGAGCGGGAAGATGTATTCTTCTCATAAG |
| | AOS30G | TCATACTTCATGTGGTAACTCCACTTCCGTCAGTCATATGACTAATATCCGCTCGACCGACATTTAGTCTGTG |
| | AOS72F | CTAGGCATATTATCTTATATGCCACAAAAAGCCGACCGTCTGGGCGTCGAGCTTGCGGGAAAGGGGTCAAC |
| | AOS73F | TCATACTTCATGTGGTAACTCCACTTCCGTCAGTCATATGACTAATGTGCGCTTTTGACGGTCTTGCCTTTTAAAC |
| | AOS01G | CTAGGCATATTATCTTATATGCCACAAAAAGCCGACCGTCTGGGCAGCGAGCAAACTTTGAGATGGCCTTGC |
| | AOS02G | TCATACTTCATGTGGTAACTCCACTTCCGTCAGTCATATGACTAATAGTCGCTCCCCTGAGACTTGGTACTGA |
| | AOS39G | CTAGGCATATTATCTTATATGCCACAAAAAGCCGACCGTCTGGGCGTGGAGCGGAACTAATTTGTATGCGAAAGTATATTG |
| | AOS40G | TCATACTTCATGTGGTAACTCCACTTCCGTCAGTCATATGACTAATTTTCGCTCAGCGGCTCCAGTGTTGC |
| | AOS74F | CTAGGCATATTATCTTATATGCCACAAAAAGCCGACCGTCTGGGCGAGGAGCTTCAGTACCAAGTCTCAGGGGAAAC |
| | AOS75F | TCATACTTCATGTGGTAACTCCACTTCCGTCAGTCATATGACTAATCTTCGCTCGGCTGTTGACCCCTTTCC |
| | AOS15G | CTAGGCATATTATCTTATATGCCACAAAAAGCCGACCGTCTGGGCGTGGAGCAAGTCCTAAGTCAACAGATCTTCTGTTG |
| | AOS16G | TCATACTTCATGTGGTAACTCCACTTCCGTCAGTCATATGACTAATATACGCTGGCTGCGTGGTTAGGACCAT |
| | AOS78F | CTAGGCATATTATCTTATATGCCACAAAAAGCCGACCGTCTGGGCTACGAGCTACCAAGTCTCAGGGGAAACTTTG |
| | AOS79F | TCATACTTCATGTGGTAACTCCACTTCCGTCAGTCATATGACTAATTCCCGCTCTGAACGGCTGTTGACCCC |
| | AOS45G | CTAGGCATATTATCTTATATGCCACAAAAAGCCGACCGTCTGGGCTTCGAGCTTTGGAGTACTCGATGGTG |
| | AOS46G | TCATACTTCATGTGGTAACTCCACTTCCGTCAGTCATATGACTAATTTCCGCTACTAATCAATATACTTTCGCATAC |
| | AOS05G | CTAGGCATATTATCTTATATGCCACAAAAAGCCGACCGTCTGGGCGAGGAGCATGGCCTTGCAAAGGGTATG |
| | AOS06G | TCATACTTCATGTGGTAACTCCACTTCCGTCAGTCATATGACTAATATACGCTCTCAAAGTTTCCCCTGAGACTTG |
| | AOS09G | CTAGGCATATTATCTTATATGCCACAAAAAGCCGACCGTCTGGGCAAGGAGCATGGTAATAAGCTGACGGACATG |
| | AOS10G | TCATACTTCATGTGGTAACTCCACTTCCGTCAGTCATATGACTAATTTTCGCTACCCTTTGCAAGGCCATCTC |
| | AOS58F | CTAGGCATATTATCTTATATGCCACAAAAAGCCGACCGTCTGGGCATGGAGCACCTAAATAGCAATATTTACC |

| | | |
|---|---|---|
| | AOS59F | TCATACTTCATGTGGTAACTCCACTTCCGTCAGTCATATGACTAATTTACGCTCAGAGTAGTGACAAGTGTTG |
| | AOS60F | CTAGGCATATTATCTTATATGCCACAAAAAGCCGACCGTCTGGGCTTGGAGCAAGTTATCAGGCATGCACCTG |
| | AOS61F | TCATACTTCATGTGGTAACTCCACTTCCGTCAGTCATATGACTAATCGACGCTTTTGACCCAAAGGTAAATATTG |
| | AOS31G | CTAGGCATATTATCTTATATGCCACAAAAAGCCGACCGTCTGGGCACGGAGCTAGTCGGACCTCTCCTTAATG |
| | AOS32G | TCATACTTCATGTGGTAACTCCACTTCCGTCAGTCATATGACTAATAGCCGCTTATCTTATGAGAAGAATACATCTTCC |
| | AOS33G | CTAGGCATATTATCTTATATGCCACAAAAAGCCGACCGTCTGGGCATCGAGCGCTAGCGGATGAAGTGATGC |
| | AOS34G | TCATACTTCATGTGGTAACTCCACTTCCGTCAGTCATATGACTAATGCCCGCTTCCCATTAAGGAGAGGTCCG |
| | AOS25G | CTAGGCATATTATCTTATATGCCACAAAAAGCCGACCGTCTGGGCGTGGAGCATGCAGTTCACAGACTAAATG |
| | AOS26G | TCATACTTCATGTGGTAACTCCACTTCCGTCAGTCATATGACTAATCTCCGCTCCATATCAACAGAAGATCTG |
| | AOS70F | CTAGGCATATTATCTTATATGCCACAAAAAGCCGACCGTCTGGGCGTCGAGCAAATTGCGGGAAAGGGGTC |
| | AOS71F | TCATACTTCATGTGGTAACTCCACTTCCGTCAGTCATATGACTAATGGGCGCTGACGGTCTTGCCTTTTAAACCG |
| | AOS64F | CTAGGCATATTATCTTATATGCCACAAAAAGCCGACCGTCTGGGCGTGGAGCTTTAAAAGGCAAGACCGTCAAATTG |
| | AOS65F | TCATACTTCATGTGGTAACTCCACTTCCGTCAGTCATATGACTAATCCTCGCTCCGATGCAATCTATTGGTTTAAAGAC |
| | AOS43G | CTAGGCATATTATCTTATATGCCACAAAAAGCCGACCGTCTGGGCGTGGAGCAGTTTTGGAGTACTCGATGG |
| | AOS44G | TCATACTTCATGTGGTAACTCCACTTCCGTCAGTCATATGACTAATCCTCGCTAATCAATATACTTTCGCATAC |
| Cloning individual Fluoride LASR variants (iPCR) | AOS55G | CGCTGCTTAGCTAATGACTCCTACCAGAATGAGCCAGATCTTCTGTTGATATGG |
| | AOS56G | TTTATGGCGAACTCCATCGCCTATAAGGACGCTTTGACTTAGGACTTGGCTG |
| | AOS57G | CGCTGCTTAGCTAATGACTCCTACCAGATCGAGCAATAGCAATATTTACCTTTG |
| | AOS58G | TTTATGGCGAACTCCATCGCCTATAACCACGCTTAGGTCAGAGTAGTGACAAG |
| | AOS59G | CGCTGCTTAGCTAATGACTCCTACCAGCATGAGCCAACAGCCGTTCAGTACCAAGTC |
| | AOS60G | TTTATGGCGAACTCCATCGCCTATAAGACCGCTACCCCTTTCCCGCAATTTG |
| | AOS61G | CGCTGCTTAGCTAATGACTCCTACCAGGGCGAGCTGATATGGATGCAGTTCAC |
| | AOS62G | TTTATGGCGAACTCCATCGCCTATAAGCGCGCTACAGAAGATCTGTTGACTTAG |
| | AOS63G | CGCTGCTTAGCTAATGACTCCTACCAGGTTGAGCTAAAAGGCAAGACCGTC |
| | AOS64G | TTTATGGCGAACTCCATCGCCTATAAGTCCGCTAACCGATGCAATCTATTG |
| | AOS65G | CGCTGCTTAGCTAATGACTCCTACCAGCATGAGCTTGCATCGGTTTAAAAGG |
| | AOS66G | TTTATGGCGAACTCCATCGCCTATAATTACGCTTCTATTGGTTTAAAGACTAGCTAC |
| | AOS67G | CGCTGCTTAGCTAATGACTCCTACCAGTTTGAGCTATGGTAATAAGCTGACGGACATG |
| | AOS68G | TTTATGGCGAACTCCATCGCCTATAATATCGCTCCCTTTGCAAGGCCATCTC |
| | AOS69G | CGCTGCTTAGCTAATGACTCCTACCAGGTTGAGCTTGAGATGGCCTTGCAAAG |

| | | |
|---|---|---|
| | AOS70G | TTTATGGCGAACTCCATCGCCTATAAGGTCGCTAGTTTCCCCTGAGACTTGGTAC |
| | AOS71G | CGCTGCTTAGCTAATGACTCCTACCAGAACGAGCCATGCACCTGGTAGCTAG |
| | AOS72G | TTTATGGCGAACTCCATCGCCTATAACGTCGCTCCTGATAACTTTTTGACCC |
| | AOS73G | CGCTGCTTAGCTAATGACTCCTACCAGGTCGAGCTAGATTGCATCGGTTTAAAAG |
| | AOS74G | TTTATGGCGAACTCCATCGCCTATAACTTCGCTTTGGTTTAAAGACTAGCTACCAG |
| | AOS75G | CGCTGCTTAGCTAATGACTCCTACCAGGCTGAGCTGCGAAAGTATATTGATTAG |
| | AOS76G | TTTATGGCGAACTCCATCGCCTATAAGTACGCTTACAAATTAGTTCCCAGC |
| | AOS77G | CGCTGCTTAGCTAATGACTCCTACCAGGTCGAGCATAGCAATATTTACCTTTGG |
| | AOS78G | TTTATGGCGAACTCCATCGCCTATAAACCCGCTTTAGGTCAGAGTAGTGACAAG |
| | AOS79G | CGCTGCTTAGCTAATGACTCCTACCAGAGTGAGCACCGTCAAATTGCGGGAAAG |
| | AOS80G | TTTATGGCGAACTCCATCGCCTATAAAGTCGCTCTTGCCTTTTAAACCGATGC |
| | AOS81G | CGCTGCTTAGCTAATGACTCCTACCAGGCTGAGCAACTAATTTGTATGCGAAAGTATATTG |
| | AOS01H | TTTATGGCGAACTCCATCGCCTATAATGCCGCTCCCAGCGGCTCCAGTG |
| | AOS02H | CGCTGCTTAGCTAATGACTCCTACCAGCCAGAGCTAAGTCAACAGATCTTCTGTTGATATG |
| | AOS03H | TTTATGGCGAACTCCATCGCCTATAACGTCGCTGGACTTGGCTGCGTGGTTAG |
| | AOS04H | CGCTGCTTAGCTAATGACTCCTACCAGGCCGAGCCCTCTCCTTAATGGGAGCT |
| | AOS05H | TTTATGGCGAACTCCATCGCCTATAACACCGCTTCCGACTATATCTTATGAGAAG |
| | AOS06H | CGCTGCTTAGCTAATGACTCCTACCAGGAGGAGCACGCAGCCAAGTCCTAAGTC |
| | AOS07H | TTTATGGCGAACTCCATCGCCTATAATGGCGCTGGTTAGGACCATGTCCGTC |
| | AOS08H | CGCTGCTTAGCTAATGACTCCTACCAGGATGAGCTCGATGGTGTTCAATGC |
| | AOS09H | TTTATGGCGAACTCCATCGCCTATAAGTACGCTGTACTCCAAAACTAATCAATATAC |
| | AOS10H | CGCTGCTTAGCTAATGACTCCTACCAGCGTGAGCAGTTTTGGAGTACTCGATGG |
| | AOS11H | TTTATGGCGAACTCCATCGCCTATAAGGCCGCTAATCAATATACTTTCGCATAC |
| | AOS12H | CGCTGCTTAGCTAATGACTCCTACCAGCTTGAGCGCAATATTTACCTTTGGGTC |
| | AOS13H | TTTATGGCGAACTCCATCGCCTATAATGCCGCTTATTTAGGTCAGAGTAGTGACAAG |
| | AOS14H | CGCTGCTTAGCTAATGACTCCTACCAGTGTGAGCTCTTCTGTTGATATGGATGC |
| | AOS15H | TTTATGGCGAACTCCATCGCCTATAATGCCGCTTCTGTTGACTTAGGACTTGG |
| | AOS16H | CGCTGCTTAGCTAATGACTCCTACCAGTCCGAGCCCAAGTCTCAGGGGAAAC |
| | AOS17H | TTTATGGCGAACTCCATCGCCTATAAGACCGCTTACTGAACGGCTGTTGACC |
| | AOS18H | CGCTGCTTAGCTAATGACTCCTACCAGTATGAGCATAGCAATATTTACCTTTGG |

| | | |
|---|---|---|
| | AOS19H | TTTATGGCGAACTCCATCGCCTATAAGGTCGCTTTAGGTCAGA GTAGTGACAAG |
| | AOS20H | CGCTGCTTAGCTAATGACTCCTACCAGCCGGAGCTAAGCTGAC GGACATGGTCC |
| | AOS21H | TTTATGGCGAACTCCATCGCCTATAAGTTCGCTTTACCATACCC TTTGCAAGG |
| | AOS22H | CGCTGCTTAGCTAATGACTCCTACCAGGGCGAGCGAGCCGCT GGGAACTAATTTG |
| | AOS23H | TTTATGGCGAACTCCATCGCCTATAAGAACGCTCAGTGTTGCAT CACTTCATCCG |
| | AOS24H | CGCTGCTTAGCTAATGACTCCTACCAGCGCGAGCATGCAGTTC ACAGACTAAATG |
| | AOS25H | TTTATGGCGAACTCCATCGCCTATAAGTACGCTCCATATCAACA GAAGATCTG |
| | AOS26H | CGCTGCTTAGCTAATGACTCCTACCAGGACGAGCGCAGCCAAG TCCTAAGTC |
| | AOS27H | TTTATGGCGAACTCCATCGCCTATAAGGGCGCTGTGGTTAGGA CCATGTCCG |
| | AOS28H | CGCTGCTTAGCTAATGACTCCTACCAGGCTGAGCTCCTTAATG GGAGCTAG |
| | AOS29H | TTTATGGCGAACTCCATCGCCTATAAAGACGCTGAGGTCCGAC TATATCTTATG |

**Supplementary Table 6. Electroporation conditions**

| Species | Potential (kV) | Current (µF) | Resistance (Ω) |
|---|---|---|---|
| *Escherichia coli* | 2.5 | 2.5 | 200 |
| *Kluyvera intermedia* | 2.5 | 2.5 | 200 |
| *Hafnia alvei* | 2.5 | 2.5 | 200 |
| *Pseudomonas chlororaphis* | 2.5 | 2.5 | 200 |
| *Comamonas testosteroni* | 1.6 | 25 | 200 |
| *Lactiplantibacillus plantarum* | 2 | 25 | 400 |
